# A molecularly defined recurrent CA3 circuit regulates network dynamics and spatial working memory

**DOI:** 10.64898/2026.10.25.761711

**Authors:** Wanpeng Cui, Haocheng Li, Wanyi Guo, Dion Y. Wang, Evan J. Bai, Yu-Qiu Jiang, Qian Sun

**Affiliations:** Department of Neurosciences, School of Medicine, Case Western Reserve University, 10900 Euclid Ave, Cleveland, OH 44106, USA

**Keywords:** hippocampus, CA3, CA1, sharp-wave ripples, memory

## Abstract

The hippocampal CA3 region supports coordinated network activity and memory processes, but how molecularly distinct CA3 neuronal populations contribute to recurrent circuit function remains poorly understood. Here we identify St18 as a molecular marker of a distinct population of deep-layer, burst-firing CA3 pyramidal neurons and generate an St18-Cre mouse line for their selective genetic targeting. In contrast to canonical CA3 pyramidal neurons, St18 neurons preferentially innervate CA3 bilaterally, with only sparse projections to CA1. Moreover, St18 neurons are highly active during endogenous CA3 sharp-wave ripples, and their optogenetic activation evokes local ripple-like activity in CA3 but not in CA1. Furthermore, chemogenetic suppression markedly reduces CA3 ripple occurrence without significantly affecting CA1 ripple occurrence and selectively impairs spatial working memory performance. Together, these findings reveal the functional significance of a molecularly defined CA3 recurrent circuit, linking neuronal heterogeneity to hippocampal network dynamics and spatial working memory.

## INTRODUCTION

The hippocampal CA3 region possesses an extensive recurrent excitatory network that plays a critical role in the neural computations underlying spatial and episodic memory^1–9^. Classically, CA3 is viewed as a relatively homogeneous auto-associative circuit that integrates cortical and dentate gyrus inputs and generates recurrent activity that is then transmitted to CA1 through the Schaffer collateral pathway^1,3–7,9–21^. However, growing evidence reveals substantial molecular, anatomical, cellular, and physiological heterogeneity among CA3 pyramidal neurons^10,22–40^. Such heterogeneity raises the question of whether molecularly distinct CA3 neuronal populations may be differentially wired into the recurrent network and make distinct contributions to hippocampal population dynamics and behavior.

One notable example of CA3 neuronal heterogeneity is recently identified “athorny” CA3 pyramidal neurons which receive minimal or very weak excitation from the dentate gyrus^26,41^. Unlike canonical CA3 pyramidal neurons, athorny neurons possess few or no thorny excrescences, preferentially reside in the deep pyramidal layer, and exhibit high-frequency burst-firing^2,26,41–50^. These distinctive features raise the possibility that athorny neurons may be uniquely integrated into the hippocampal network to support specific neural computations^2,3,43^. Previous studies have implicated these neurons in coordinated CA3 population activity, including sharp-wave ripples (SWRs)^2,26^, a prominent form of synchronized hippocampal activity associated with memory processing^51–68^. However, the molecular identity of athorny neurons, their functional circuit architecture, how they interact within local and downstream hippocampal circuits, and their causal contributions to network activity and behavior remain poorly understood.

Here, we identify Suppression of Tumorigenicity 18 (St18) as a molecular marker of deep-layer, burst-firing CA3 athorny neurons and have generated a knock-in St18-P2A-iCre mouse line that enables their selective targeting and manipulation. We demonstrate that St18 neurons exhibit a distinctive output architecture, preferentially innervating the CA3 recurrent network while providing only sparse projections to CA1, which contrasts sharply with canonical CA3 pyramidal neurons. Within CA3, these neurons strongly activate local inhibitory circuits and are highly active during spontaneous SWRs. In vivo recordings and cell-type-specific manipulations further demonstrate that St18 neurons contribute to local CA3 ripple dynamics, while their suppression selectively impairs spatial working memory. Together, these findings uncover a molecularly defined CA3 circuit that selectively regulates local hippocampal network dynamics and spatial working memory.

## RESULTS

### Single-cell transcriptomics identifies St18 as a marker of deep-layer CA3 glutamatergic neurons

To identify molecularly distinct CA3 pyramidal neuron populations, we analyzed single-cell RNA-sequencing data from the Allen Institute^69^. After selecting hippocampal CA3 glutamatergic neurons (n = 1,822), Uniform manifold approximation and projection (UMAP) analysis resolved the population into two major transcriptional domains comprising eight subclusters, revealing remarkable molecular heterogeneity among CA3 pyramidal neurons (Fig. 1A-B). All subclusters expressed the pan-neuronal marker *Snap25*, the glutamatergic marker *Slc17a7* (*Vglut1*), the pyramidal neuron marker *Camk2a*, and the CA3-enriched marker *Grik4*, while lacking the inhibitory neuronal marker *Slc32a1* (*Vgat*), confirming their identity as excitatory CA3 pyramidal neurons (Fig. 1C; fig. S1A).

**Figure 1.**
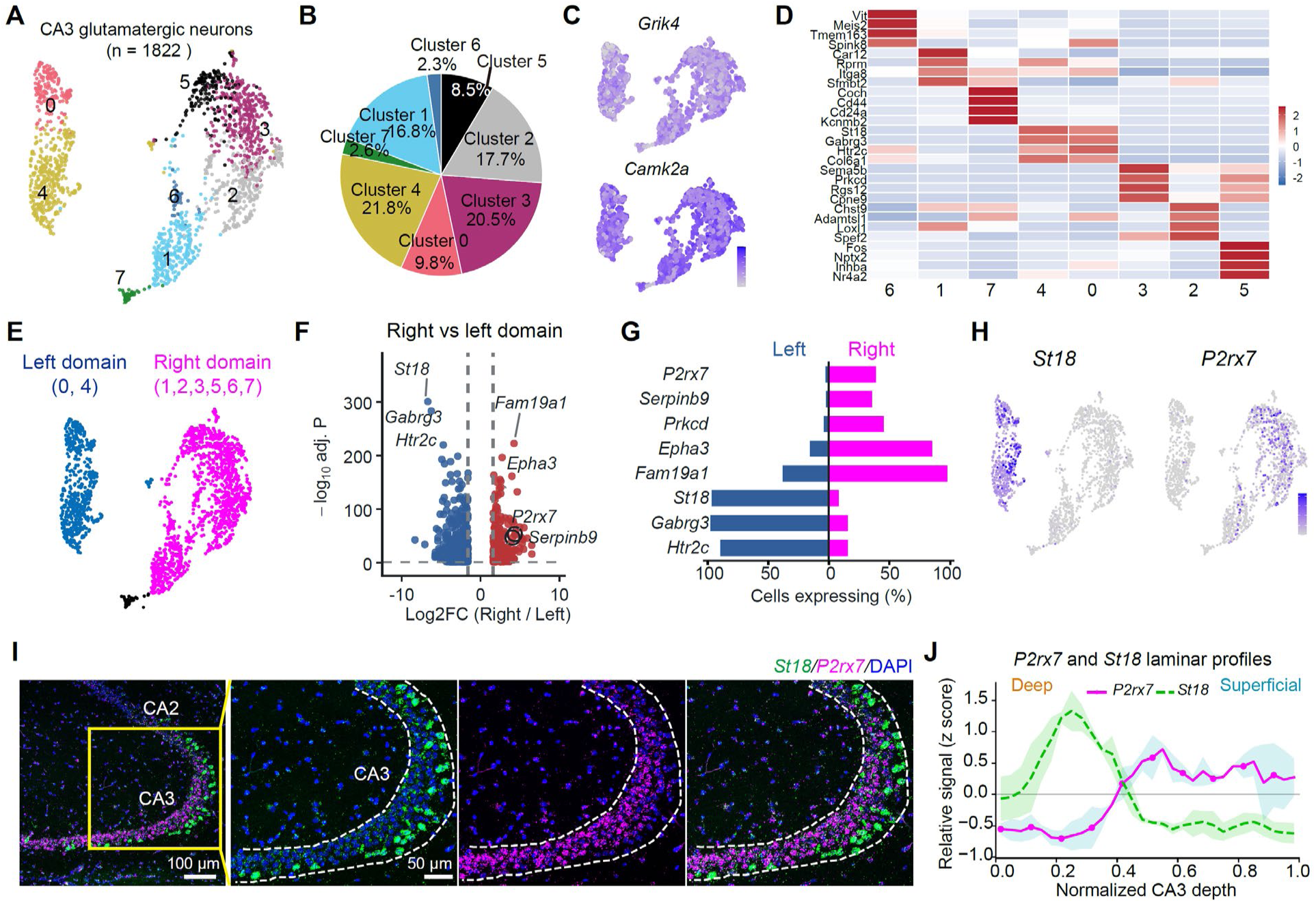
Single-cell transcriptomic analysis identifies *St18* as a marker of deep-layer CA3 pyramidal neurons. (**A**) UMAP representation of 1,822 mouse CA3 glutamatergic pyramidal neurons, resolving eight transcriptional subclusters. (**B**) Pie chart showing percentage of cells of each subclusters among the CA3 population. (**C**) UMAP feature plots showing expression of the CA3-enriched marker *Grik4* and the pyramidal neuron marker *Camk2a*. Expression scale, 0-6. (**D**) Heatmap showing the top four marker genes for each CA3 subcluster. (**E**) UMAP representation of the two major CA3 transcriptional domains. Clusters 0 and 4 were grouped as the left domain (dark blue), whereas clusters 1, 2, 3, 5, and 6 were grouped as the right domain (magenta). Cluster 7, which occupied an intermediate transcriptional position, is shown in black. (**F**) Volcano plot showing differentially expressed genes between the left and right domains. Colored points indicate genes with Bonferroni-adjusted *p* < 0.05 and expression difference greater than threefold. Dashed lines indicate thresholds. (**G**) Percentage of cells within the left and right domains expressing the indicated candidate marker genes. (**H**) UMAP feature plots showing the expression distributions of *St18* and *P2rx7* across CA3 pyramidal neurons. Scale, [0 4]. (**I**) Representative RNAscope images showing the laminar distribution of *St18* and *P2rx7* within CA3. (**J**) Quantification of *St18* and *P2rx7* expression across the deep-to-superficial axis of the CA3 pyramidal cell layer. Signals were normalized as z-scores and smoothed using a centered three-bin moving average. Solid lines, mean ± SEM, n = 2 mice.

Differential gene expression analysis identified several genes enriched in the two transcriptional domains. Among these, *St18* exhibited the highest prevalence and specificity for one domain, with *Htr2c* and *Gabrg3* also enriched in the same population, whereas *P2rx7* preferentially marked the opposing domain (Fig. 1D-F; fig. S1B). We next examined the spatial distribution of these candidate markers using the Allen Brain Atlas in situ hybridization database. *St18* displayed a highly restricted expression pattern concentrated in the deep portion of the CA3 pyramidal layer, near the stratum pyramidale/stratum oriens border (fig. S1D), supporting *St18* as a candidate marker of the transcriptionally distinct deep-layer CA3 population.

We further validated the laminar expression patterns of *St18* and *P2rx7* using RNAscope. *St18* expression peaked in the deep CA3 pyramidal layer, whereas *P2rx7* expression increased toward the middle-to-superficial layer, with limited overlap between the two markers (Fig. 1H-J; fig. S1C). Thus, the two transcriptional domains identified by single-cell analysis correspond to spatially distinct populations along the deep-to-superficial axis of CA3, with *St18* preferentially marking deep-layer CA3 pyramidal neurons. In addition, single-nucleus transcriptomic analysis of human hippocampus revealed enrichment of *ST18* in a distinct cluster of human CA3 glutamatergic neurons^70^ (fig. S1E-F), suggesting that an *ST18*-enriched CA3 neuronal population may also be present in humans^50^. Finally, co-staining for *St18* and *Slc17a7* revealed that *St18*-expressing neurons accounted for 8.24 ± 0.76% and 13.95 ± 1.91% of glutamatergic neurons in dorsal and ventral CA3, respectively, and the proportion of *St18*-expressing neurons was higher in distal (CA3a: 10.03 ± 2.34%, CA3b: 11.17 ± 1.77%) than in proximal CA3 (CA3c: 2.76 ± 0.71%, *p* = 0.01, one-way ANOVA, fig. S2).

Together, these analyses identify *St18* as a molecular marker of a spatially distinct deep-layer CA3 pyramidal neuron population, providing a means to genetically access this population.

### St18 defines athorny CA3 pyramidal neurons

We asked whether St18 labels CA3 neurons with the characteristic features of the previously described athorny neurons^2,26,41,43,46^. To test this, we generated a St18-P2A-iCre (St18-Cre) knock-in mouse line and selectively labeled St18 neurons by injecting Cre-dependent AAV-DIO-EYFP into CA3 (Fig. 2A). EYFP-labeled neurons were concentrated in the deep CA3 pyramidal layer, near the stratum pyramidale/stratum oriens border, particularly in distal CA3 (Fig. 2B,D). RNAscope confirmed that 98.4% of EYFP-labeled neurons expressed endogenous *St18* (362/368 cells, 2 mice, Fig. 2C), demonstrating faithful recapitulation of endogenous gene expression by the knock-in line. Unlike canonical thorny CA3 pyramidal neurons, these cells possessed few or no thorny excrescences (Fig. 2E-G).

**Figure 2.**
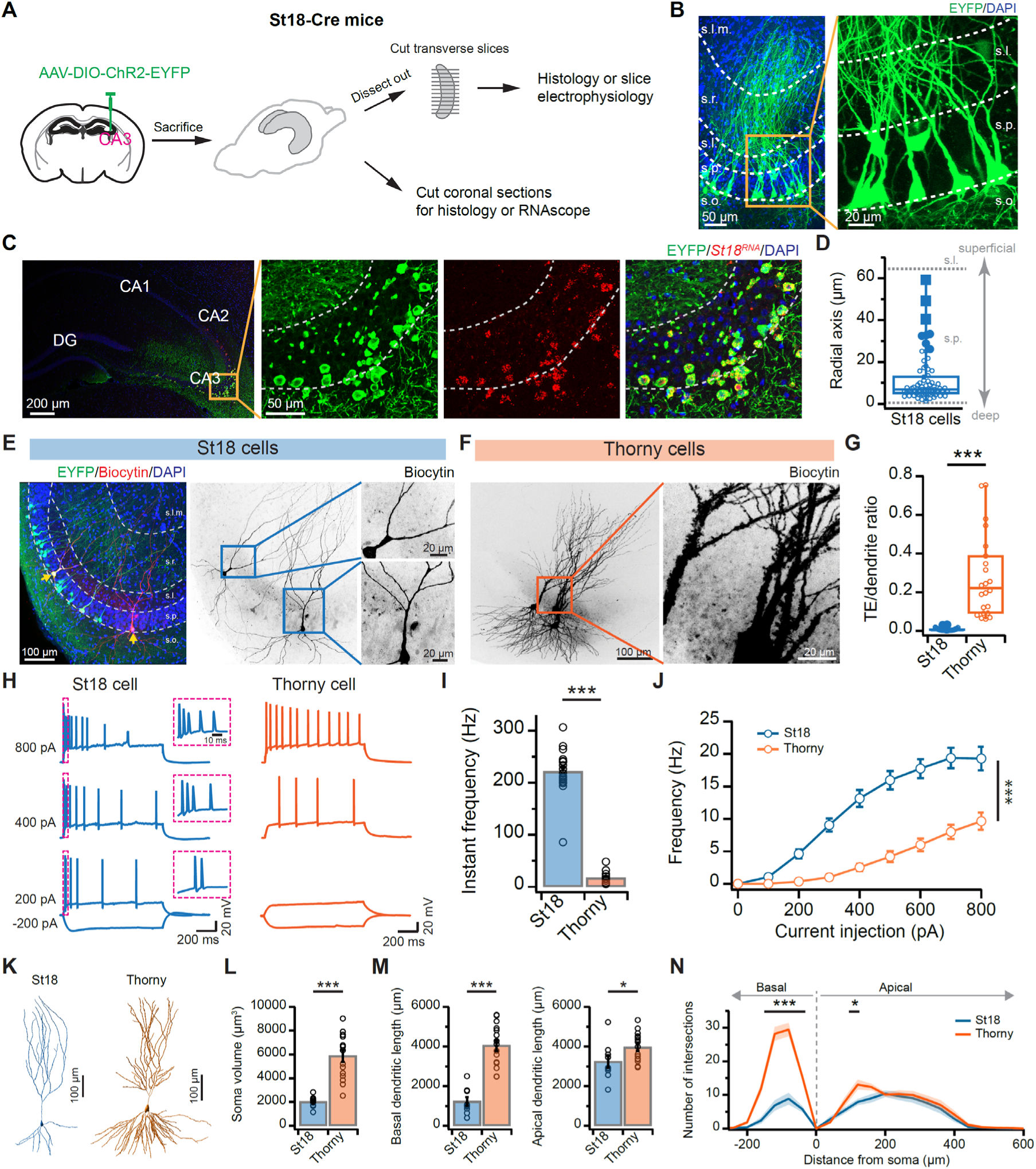
St18 molecularly defines deep-layer, burst-firing CA3 pyramidal neurons. **(A)** Generation of St18-Cre mouse line and experimental procedure. Top, Schematic of the *St18* gene structure and the Cre recombinase knock-in strategy. A P2A-iCre cassette was inserted immediately downstream of exon 24 of the *St18* locus to enable Cre expression in St18-expressing cells. Bottom, AAV-DIO-EYFP was injected into the CA3 region of St18-Cre mice for histological analysis, RNAscope, and slice electrophysiology. **(B)** Confocal image showing EYFP-labeled St18 neurons in CA3. **(C)** Confocal images showing a high level of coexpression of endogenous *St18* RNA and EYFP (362/368 cells/2 mice). **(D)** Radial distribution of St18 neurons within the CA3 pyramidal cell layer (*n* = 72 cells). Filled symbols represent outliers. **(E)** Representative confocal images of biocytin-filled St18 neurons following whole-cell recording. Enlarged views show the absence or sparse presence of thorny excrescences. **(F)** Representative biocytin-filled canonical CA3 pyramidal neurons. Enlarged views show abundant thorny excrescences. **(G)** Quantification of ratio of thorny excrescences relative to corresponding dendritic volume in St18 (*n* = 28) and canonical thorny CA3 pyramidal neurons (*n* = 22). **(H)** Representative current-clamp recordings showing firing responses of St18 and thorny CA3 neurons during 1-s depolarizing current injections. **(I)** Initial instantaneous firing frequency of St18 (*n* = 21) and thorny neurons (*n* = 12), calculated from the first two action potentials evoked by an 800-pA current injection. \*\*\**p* < 0.001, unpaired *t* test. **(J)** Frequency-current (F-I) relationship for St18 (*n* = 21) and thorny neurons (*n* = 11). \*\*\**p* < 0.001, repeated-measures two-way ANOVA. **(K)** Representative three-dimensional reconstructions of St18 and thorny neurons generated with Imaris software. **(L)** Soma volume of St18 (*n* = 9) and thorny neurons (*n* = 15). Mean ± SEM. \*\*\**p* < 0.001, unpaired *t* test. **(M)** Quantification of basal dendritic length (left) and apical dendritic length (right) in St18 (*n* = 9) and thorny neurons (*n* = 15). Mean ± SEM. \**p* < 0.05, \*\*\**p* < 0.001, unpaired *t* test. **(N)** Sholl analysis of basal and apical dendritic arborization in St18 (*n* = 9) and thorny neurons (*n* = 15). Solid lines indicate mean and shaded areas indicate SEM. \**p* < 0.05, \*\*\**p* < 0.001, repeated-measures two-way ANOVA with Tukey’s multiple-comparisons test.

Whole-cell current-clamp recordings revealed that St18 neurons exhibited robust high-frequency burst-firing in response to somatic current injection (initial instantaneous frequency induced by 800-pA injection, 221.7 ± 9.1 Hz; *n* = 21 cells; Fig. 2E,H,I). In contrast, canonical thorny neurons (confirmed by post hoc biocytin staining) displayed regular-spiking activity with a substantially lower initial instantaneous firing frequency (17.4 ± 3.6 Hz; *n* = 12 cells; *p* = 7.9 × 10^-17^, unpaired *t* test; Fig. 2F,H,I) and generated fewer action potentials in response to identical current injections (Fig. 2J). Notably, the distributions of initial firing frequency did not overlap between the two populations (Fig. 2I), indicating a clear physiological distinction. Moreover, St18 neurons (*n* = 9 cells) possessed significantly smaller somata, shorter apical and basal dendrites, fewer branch intersections in basal dendrites than thorny neurons (*n* = 15 cells), closely resembling the morphology previously described for athorny CA3 neurons (Fig. 2K-N).

Together, these findings demonstrate that St18 neurons constitute a distinct CA3 pyramidal neuron subtype defined by their deep-layer location, paucity of thorny excrescences, distinct dendritic architecture, and intrinsic burst-firing properties. These defining anatomical, morphological, and electrophysiological characteristics closely match those of the previously described athorny CA3 pyramidal neurons^26^, providing a molecular identity for this neuronal population.

### St18 neurons preferentially innervate CA3 and project sparsely to CA1

To map St18 cell outputs, we unilaterally injected Cre-dependent AAV-ChR2-EYFP into CA3 of St18-Cre mice and inspected the expression of EYFP in transverse sections from dissected hippocampi. Strikingly, EYFP-labeled axons were largely confined to CA3 in both hemispheres, with only sparse projections to CA1 (Fig. 3A-B, fig. S3). Quantification of contralateral axonal fluorescence in the dorsal hippocampus, where somatodendritic fluorescence was less likely to confound axonal measurements, revealed dense innervation of the stratum oriens in distal CA3 but little labeling in proximal CA3 or other CA3 layers (Fig. 3A,B,E,F). This projection pattern differed markedly from that of canonical CA3 pyramidal neurons labeled in Grik4-Cre mice, whose axons form both recurrent collaterals within CA3 and Schaffer collaterals to CA1 (Fig. 3C-F). In the ipsilateral ventral hippocampus, St18 axons shifted toward the stratum radiatum in CA3 while maintaining minimal projections to CA1 (Fig. 3G-J), suggesting that sparse CA1 innervation is maintained across the longitudinal levels. A similar topographic pattern of St18 projections was observed in coronal sections (fig. S3).

**Figure 3.**
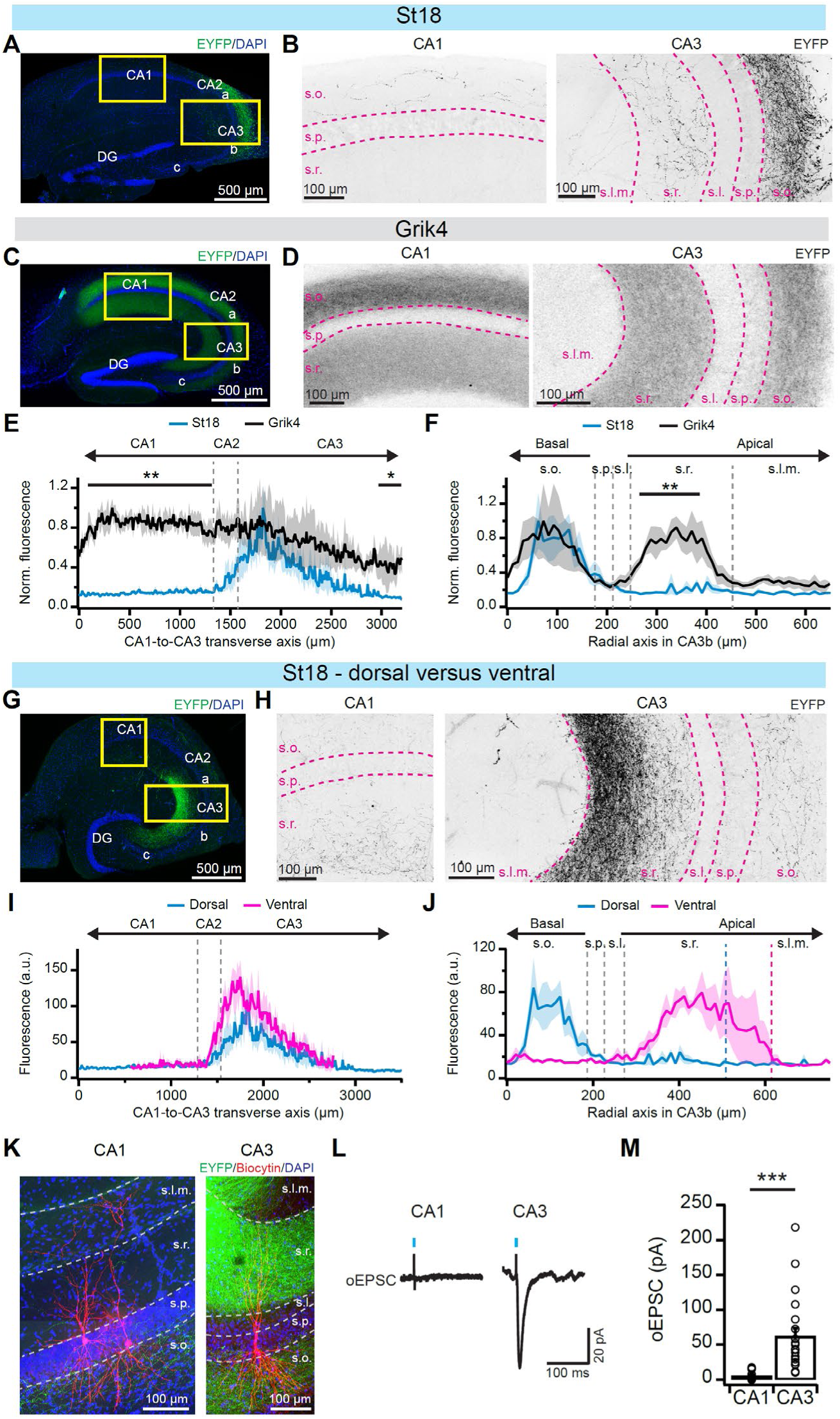
St18 neurons preferentially innervate CA3 while largely avoiding CA1. **(A)** Confocal images showing EYFP expression in the contralateral dorsal hippocampus of St18-Cre mice following unilateral injection of AAV-DIO-EYFP-ChR2 into CA3. **(B)** Higher-magnification views of the boxed regions in CA1 and CA3 shown in (A). EYFP signal is pseudo-colored black. **(C)** Confocal images showing EYFP expression in the contralateral dorsal hippocampus of Grik4-Cre mice following unilateral injection of AAV-DIO-EYFP-ChR2 into CA3. **(D)** Higher-magnification views of the boxed regions in CA1 and CA3 shown in (C). EYFP signal is pseudo-colored black. **(E)** Quantification of normalized EYFP fluorescence intensity along the CA1-to-CA3 transverse axis in St18 (*n* = 4) and Grik4 (*n* = 4) mice. Fluorescence was measured in stratum oriens and normalized to the peak fluorescence intensity for each experiment. Solid lines and shaded areas represent mean ± SEM. \**p* < 0.05, \*\**p* < 0.01, repeated-measures two-way ANOVA with Tukey’s multiple-comparisons test. **(F)** Quantification of normalized EYFP fluorescence intensity along the radial axis of CA3b in St18 (*n* = 4) and Grik4 (*n* = 4) mice. Fluorescence was measured in stratum oriens and normalized to the peak fluorescence intensity for each experiment. Solid lines and shaded areas represent mean ± SEM. \*\**p* < 0.01, repeated-measures two-way ANOVA with Tukey’s multiple-comparisons test. **(G)** Confocal images showing EYFP expression in the ipsilateral ventral hippocampus of St18-Cre mice following unilateral injection of AAV-DIO-EYFP-ChR2 into CA3. **(H)** Higher-magnification views of the boxed regions in CA1 and CA3 shown in (G). EYFP signal is pseudo-colored black. **(I)** Quantification of normalized EYFP fluorescence intensity along the transverse CA1-to-CA3 axis in the dorsal (*n* = 4) and ventral (*n* = 5) hippocampus of St18-Cre mice. Fluorescence was measured in stratum oriens for the dorsal hippocampus and stratum radiatum for the ventral hippocampus. Solid lines and shaded areas represent mean ± SEM. **(J)** Quantification of normalized EYFP fluorescence intensity along the radial axis of CA3b in the dorsal (*n* = 4) and ventral (*n* = 5) hippocampus of St18-Cre mice. **(K)** Representative confocal images of biocytin-filled CA1 (left) and CA3 (right) pyramidal neurons following whole-cell recordings. **(L, M)** Representative voltage-clamp recordings of optogenetically evoked EPSCs (oEPSCs) in CA1 and CA3 pyramidal neurons (L) and summary data (M). CA1, *n* = 15 cells; CA3, *n* = 20 cells. \*\*\**p* < 0.001, unpaired *t* test.

We next examined the functional connectivity of these projections using ChR2-assisted whole-cell recordings. Photostimulation of St18 axons reliably evoked excitatory postsynaptic responses in CA3 pyramidal neurons (oEPSC = 61.38 ± 12.39 pA, *n* = 20), whereas CA1 pyramidal neurons exhibited little or no excitation (oEPSC = 5.03 ± 1.58 pA, *n* = 15, *p* = 0.00044, unpaired *t* test, Fig. 3K-M), consistent with the anatomical projection pattern. None of the recorded CA3 pyramidal neurons fired action potentials in response to light stimulation (0/20 cells).

Together, these findings identify St18 neurons as a distinct CA3 excitatory neuronal subtype for recurrent connectivity, with projections confined within CA3 and sparse output to CA1. This projection pattern suggests that St18 neurons preferentially influence local CA3 processing rather than directly transmitting excitatory output to CA1.

### St18 neurons strongly activate disynaptic inhibition in the CA3 network

We next examined how the local CA3 circuit responded to St18 neuron activation. Whole-cell recordings from CA3 interneurons revealed that St18 neurons strongly excited local inhibitory circuits (Fig. 4A-F). Light-evoked EPSCs were substantially larger in CA3 interneurons than in CA3 pyramidal neurons, consistent with stronger functional excitation of the inhibitory neurons (Fig. 4C). Repetitive light stimulation at 20 Hz resulted in sustained firing of these interneurons (Fig. 4D-E). Overall, light stimulation of St18 axons elicited excitatory responses in 95% of randomly sampled CA3 interneurons (19/20 cells), and 80% of interneurons (16/20 cells) reached action potential threshold (Fig. 4F).

**Figure 4.**
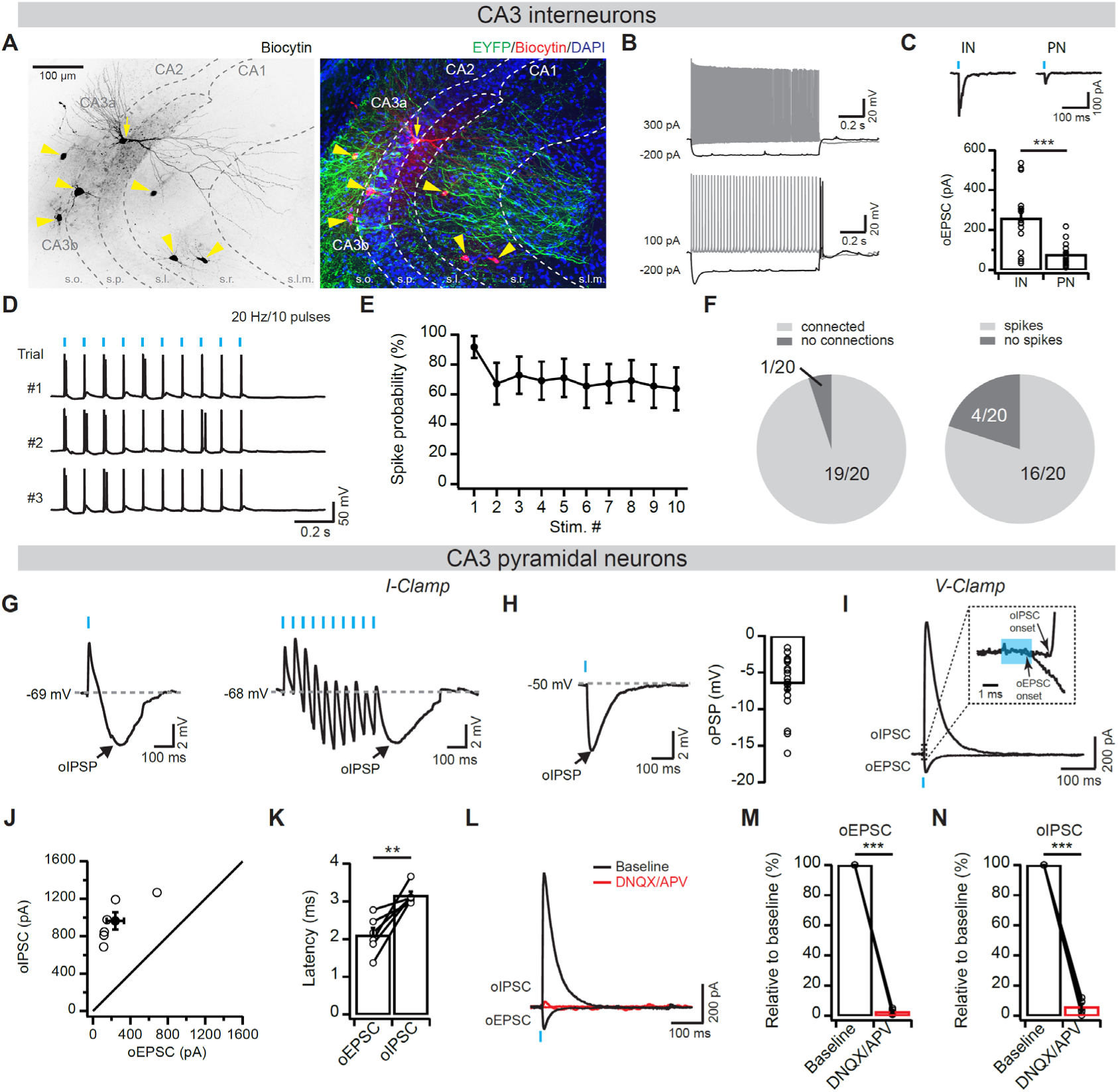
St18 neurons strongly excite inhibitory circuits in CA3. (**A**) Confocal images of biocytin-filled CA3 pyramidal neurons (arrow) and interneurons (arrowhead) that underwent whole-cell recordings. St18-Cre mice were injected with AAV-DIO-ChR2. (**B**) Representative voltage traces in response to the indicated current injections from two CA3 interneurons. (**C**) Top, representative oEPSC traces recorded from a CA3 interneuron (left) and a CA3 pyramidal neuron (right). Bottom, group data comparing oEPSC amplitudes between CA3 interneurons (*n* = 16 cells) and pyramidal neurons (*n* = 14 cells). (**D**) Representative voltage traces showing action potentials evoked by repetitive light stimulation in a CA3 interneuron. (**E**) Quantification of spike probability during light stimulation (*n* = 11 cells). (**F**) Pie charts showing the proportion of CA3 interneurons exhibiting oEPSCs (left) or light-evoked spikes (right). Numbers indicate the number of cells. (**G**) Optically evoked biphasic changes in membrane potential at the resting potential recorded in current clamp. Left and right show responses to a single light pulse and a train of 20-Hz light pulses, respectively. Note the large hyperpolarization (oIPSP). (**H**) Same as in (**A**), except that the membrane potential was depolarized to −50 mV to increase the driving force for inhibition. Left, representative trace; right, group data (*n* = 19 cells). (**I**) Optically evoked EPSCs (oEPSCs; holding potential = −70 mV) and IPSCs (oIPSCs; holding potential = 0 mV) recorded in voltage clamp using cesium-based internal solutions. (**J**) oIPSC amplitude plotted as a function of oEPSC amplitude. Open and filled symbols represent individual cells and group means, respectively. (**K**) Group data comparing the onset latencies of oEPSCs and oIPSCs (*n* = 6 cells). (**L–N**) Representative traces of oEPSCs and oIPSCs in the absence and presence of DNQX/APV (**L**), and group data for oEPSC (**M**, *n* = 6 cells) and oIPSC amplitudes (**N**, *n* = 6 cells). \*\**p* < 0.05, \*\*\**p* < 0.001. Error bars represent SEM.

Consistent with strong interneuron excitation, activation of St18 neurons generated robust hyperpolarizing responses in CA3 pyramidal neurons under current-clamp conditions (Fig. 4G). These inhibitory responses were enhanced when neurons were depolarized to −50 mV to increase the driving force for chloride-mediated inhibition (*n* = 19 cells, Fig. 4H). To distinguish direct excitation from recruited inhibition, we performed voltage-clamp recordings from the same CA3 pyramidal neurons using cesium-based internal solutions (Fig. 4I). Stimulation of St18 axons evoked both oEPSCs at −70 mV and oIPSCs at 0 mV, with an inhibition-to-excitation ratio of 5.53 ± 0.80 (*n* = 6 cells, Fig. 4J). oIPSCs exhibited significantly delayed onset relative to oEPSCs (Fig. 4K) and blockade of AMPA/NMDA receptors with DNQX/APV abolished both oEPSCs and oIPSCs, consistent with St18-driven inhibition arising through glutamatergic excitation of local interneurons (Fig. 4L-N).

Together, these findings demonstrate that St18 neurons strongly excite CA3 inhibitory circuits and impose strong disynaptic inhibition onto CA3 pyramidal neurons, revealing a mechanism by which a recurrent-projecting CA3 subtype regulates local network activity.

### Activation of St18 neurons evokes CA3 but not CA1 ripple-like activity

To determine whether St18 neurons participate in hippocampal network dynamics in vivo, we performed optotagged single-unit recordings from CA3 neurons expressing ChR2 in St18-Cre mice under light anesthesia (Fig. 5A). Among 122 isolated CA3 units, 12 met stringent optotagging criteria based on short-latency, reliable responses to 100-ms light pulses, identifying putative optotagged St18 neurons (Fig. 5B). These neurons exhibited highly time-locked responses to optical stimulation, with a representative cell showing a median first-spike latency of 5.45 ms, a first-spike jitter of 1.22 ms, and 91.7% response reliability. Optotagged St18 neurons exhibited high light-response reliability across recorded units.

**Figure 5.**
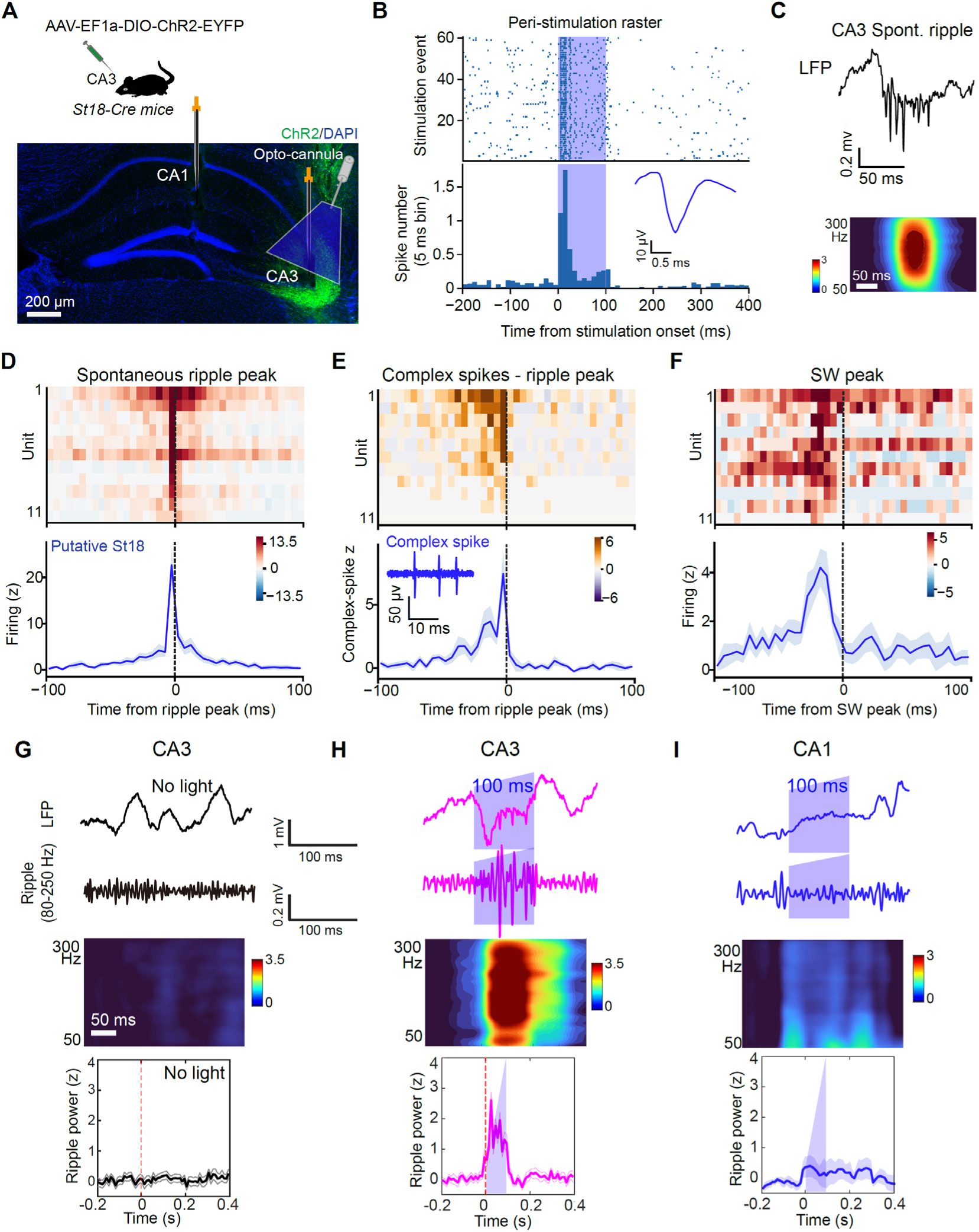
St18 neurons promote ripple-like activity in CA3 but not CA1. (**A**) Experimental schematic for optogenetic tagging and in vivo electrophysiological recordings. AAV-DIO-ChR2-EYFP was injected into CA3 of St18-Cre mice. Carbon-fiber electrodes were positioned in either CA3 or CA1 for LFP and unit recordings, and an optical fiber was placed above CA3 for photo-stimulation. Bottom, representative ChR2-EYFP expression in CA3. (**B**) Representative peri-stimulus raster plot and peri-event time histogram from a putative St18 neuron tagged by ChR2. Blue shading indicates the photo-stimulation period. Lower inset, representative extracellular spike waveform from the illustrated unit. (**C**) Representative LFP traces showing spontaneous CA3 ripple event and averaged time frequency decomposition. The time-frequency analysis includes 1,020 spontaneous events (17 recording sessions/6 mice). Color scale, 0-3. (**D**) Firing of putative St18 neurons aligned to the peak of spontaneous ripple events. Top, heatmap showing the z-scored firing rate of individual St18 units aligned to ripple peaks. Color scale, −13.5 to 13.5. Bottom, population-averaged firing rate aligned to the ripple peak. (**E**) Complex-spike activity of putative St18 neurons aligned to spontaneous ripple peaks. Top, heatmap showing complex-spike activity of individual St18 units aligned to ripple peaks. Color scale, −6 to 6. Bottom, population-averaged complex-spike activity aligned to the ripple peak. Inset, representative complex-spike waveform. (**F**) Activity of putative St18 neurons aligned to spontaneous CA3 sharp-wave peaks. Top, heatmap showing the firing activity of individual St18 units aligned to sharp-wave peaks. Color scale, −5 to 5. Bottom, population-averaged firing rate aligned to the sharp-wave peak. Solid line and shading, mean ± SEM. *n* = 11 units from 6 mice. (**G**) Control recordings from CA3 without optical stimulation. Top, representative broadband LFP and ripple-band-filtered traces during photo-stimulation. Blue shading indicates the stimulation period. Middle, average time-frequency representation over 50-300 Hz. Color scale, 0-3.5. Bottom, peri-stimulus ripple-band power. *n* = 210 stimulation trials from 3 mice. (**H**) Optogenetic activation of St18 neurons evokes local ripple-like activity in CA3. Top, representative broadband LFP and ripple-band-filtered traces during photo-stimulation. Blue shading indicates the stimulation period. Middle, average time-frequency representation aligned to photo-stimulation. Color scale, 0-3.5. Bottom, peri-stimulus ripple-band power. Solid line and shading indicate mean ± SEM. *n* = 232 stimulation trials from 4 mice. (**I**) CA1 recording of LFP and ripple-like events in CA3 St18 activation. Top, representative CA1b broadband LFP and ripple-band-filtered traces during CA3 St18-neuron photo-stimulation. Blue shading indicates the stimulation period. Middle, average time-frequency representation aligned to photo-stimulation. Color scale, 0-3. Bottom, peri-stimulus ripple-band power. Solid line and shading indicate mean ± SEM. *n* = 240 stimulation trials from 4 mice.

We next examined St18 neuronal activity during spontaneous CA3 SWR events. Eleven of 12 optotagged St18 neurons showed a marked increase in firing around spontaneous ripple events (Fig. 5C-D). St18 neurons also exhibited an increase in complex-spike firing during ripples, consistent with strong activity of this population during high-frequency CA3 network activity (Fig. 5E). Notably, St18 firing increased before the peak of spontaneous SWR events (Fig. 5F), indicating that St18 neurons are activated around the onset of synchronized CA3 population activity. Together, these observations suggest that St18 neurons contribute to the temporal dynamics of endogenous CA3 SWRs.

We next asked whether selective St18 activation can evoke local ripple-like activity. We optogenetically stimulated St18 neurons while simultaneously recording local field potentials (LFPs) in CA3 or downstream CA1^55,71,72^. In ChR2, but not control mice, brief optical stimulation reliably evoked high-frequency ripple-like activity in CA3, with ripple power increasing during the 100-200-ms stimulation period (Fig. 5G-H; fig. S4). In contrast, the same stimulation protocol did not produce detectable ripple-like responses in CA1 (Fig. 5I). We next performed closed-loop stimulation to test whether activation of St18 neurons specifically during ongoing CA1 SWRs could enhance SWRs. Light pulses were delivered to stimulate St18 neurons in CA3 after SWRs were detected in CA1 ∼20 ms after ripple onset (fig. S5A). We found nearly identical ripple powers and durations between light and no-light conditions (fig. S5B-E). Thus, St18 neuron activation evokes local CA3 ripple-like activity without a detectable corresponding response in CA1.

Overall, these findings demonstrate that St18 neurons are highly active during endogenous CA3 SWRs and that their selective activation can evoke local ripple-like activity. The predominantly local response is consistent with the preferential CA3 innervation of St18 neurons and supports a role for these cells in regulating local recurrent network activity.

### Suppression of St18 neurons reduces CA3 but not CA1 ripple occurrence

We next asked whether St18 neuron activity supports endogenous ripple activity. We bilaterally expressed the inhibitory DREADD hM4Di in St18 neurons using a Cre-dependent viral strategy (AAV-hSyn-DIO-hM4Di-mCherry; Fig. 6A and fig. S6A), with AAV-DIO-mCherry serving as the control. We then recorded LFPs from CA3 and CA1 in lightly anesthetized mice following saline or clozapine-N-oxide (CNO) administration in a within-animal design.

**Figure 6.**
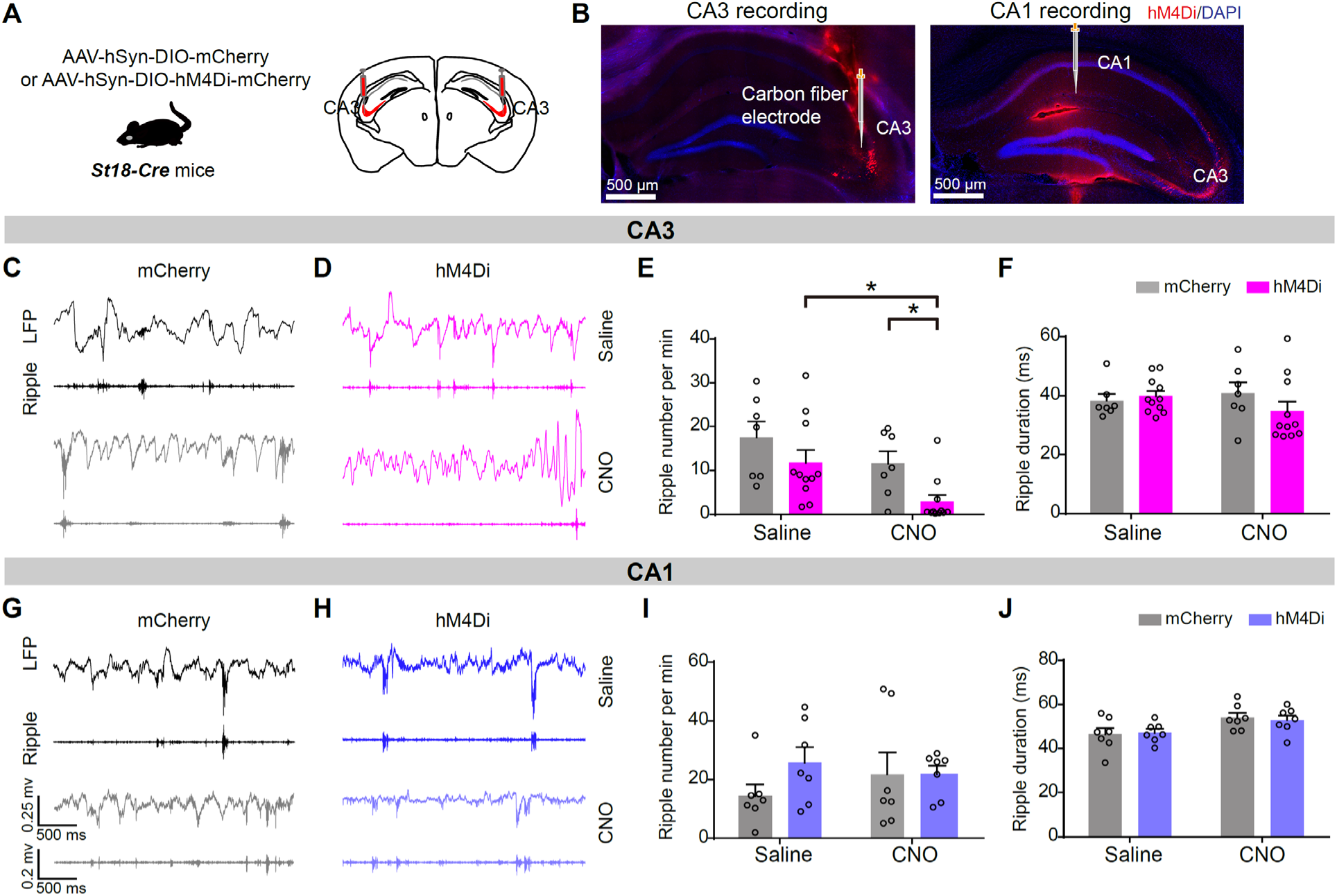
Suppression of St18 neurons reduces CA3 ripple activity. (**A**) Schematic showing Cre-dependent expression of inhibitory hM4Di-mCherry in CA3 St18 neurons of St18-Cre mice following injection of AAV-hSyn-DIO-hM4Di-mCherry. Control mice received AAV-DIO-mCherry. (**B**) Representative histological verification of viral expression and recording sites in CA3 and CA1. (**C-D**) Representative raw CA3 LFP traces and ripple-band-filtered traces from mCherry and hM4Di-expressing mice and after saline or CNO treatment. Black, saline and CNO traces of mCherry group, magenta, hM4Di. (**E**) Quantification of CA3 ripple rate. Two-way ANOVA, (F(1,16) = 7.52, *p* = 0.015). hM4Di, Saline vs CNO, *p* = 0.035, mCherry, saline vs CNO, *p* = 0.33. (**F**) Quantification of CA3 ripple duration. No significant main effect of treatment was observed. Two-way ANOVA, F(1,16) = 0.19, *p* = 0.67. Cherry, Saline vs CNO, *p* = 0.84; hM4Di, saline vs CNO, *p* = 0.34. *n* = 7 mCherry mice and 11 hM4Di mice. (**G-H**) Representative CA1 LFP and ripple-band-filtered traces following saline or CNO administration in mCherry mice (G) and hM4Di-expressing mice (H). (**I**) Quantification of CA1 ripple rate. Two-way ANOVA, F(1,12) = 0.13, *p* = 0.72), mCherry, saline vs CNO, *p* = 0.49; and hM4Di, saline vs CNO, *p* = 0.81. (**J**) Quantification of CA1 ripple duration. Two-way ANOVA, F(1,12) = 6.67, *p* = 0.02, mCherry, saline vs CNO, *p* = 0.12 or hM4Di, saline vs CNO, *p* = 0.26. *n* = 7 mice per group. Data are presented as mean ± SEM. Post hoc comparisons were Sidak’s multiple comparisons test. \**p* < 0.05. NS, not significant.

Control mice exhibited robust ripple activity after both saline and CNO administration. In contrast, CNO significantly reduced ripple occurrence in hM4Di mice, from 9.2 to 1.2 events per minute (*p* = 0.035; Fig. 6C-E, CNO, 2 mg/kg, i.p.). Ripple duration was not altered (Fig. 6F). This effect was region-specific: suppression of St18 neurons did not significantly alter ripple occurrence or duration in CA1 (Fig. 6G-J). Because ripple events preferentially occur during non-theta states, we asked whether the reduction in CA3 ripple occurrence could instead reflect a shift in network state. Restricting the analysis to non-theta periods yielded a similar reduction in CA3 ripple occurrence following St18 neuron suppression (fig. S6B). Likewise, no significant changes in non-theta ripple dynamics were detected in CA1 (fig. S6C).

Together, these findings show that suppression of St18 neurons selectively reduces the occurrence of endogenous ripple events in CA3 without altering ripple duration. The absence of a statistically significant CA1 effect is consistent with a preferential contribution of St18 neurons to local CA3 ripple dynamics, although other effects on CA1 cannot be excluded.

### St18 neuron suppression impairs performance in spatial working memory tasks

We next examined the behavioral consequences of suppressing St18 neurons. Chemogenetic inhibition of St18 neurons did not alter locomotor or exploratory activity in the open field test or performance in the novel object recognition task, suggesting that suppression did not produce broad behavioral or recognition memory deficits (Fig. 7A-E, fig. S7A-C). Next, we performed contextual fear conditioning to examine whether St18 neurons participate in CA3-dependent associative long-term memory. Retrieval of recent and remote fear memory was tested 24 hours and 20 days after foot shocks, respectively. Interestingly, suppression of St18 neurons before retrieval did not alter freezing during either recent or remote memory retrieval (Fig. 7F-G), suggesting that St18 suppression does not impair contextual fear memory retrieval under these conditions.

**Figure 7.**
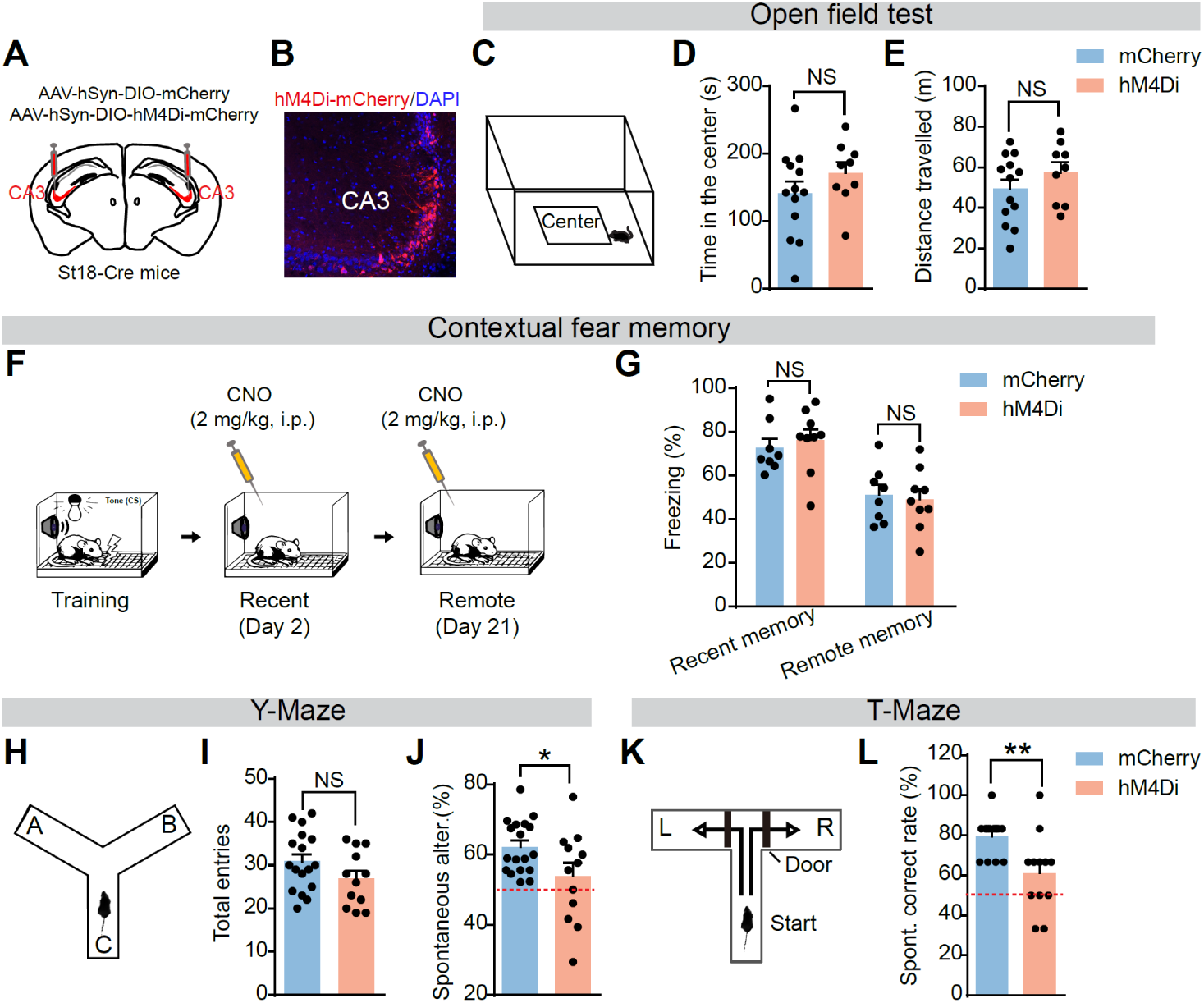
Chemogenetic suppression of St18 neurons impairs spatial working memory. (**A**) Experimental schematic showing bilateral injection of AAV-hSyn-DIO-hM4Di-mCherry or AAV-hSyn-DIO-mCherry into CA3 of St18-Cre mice. CNO (2.0 mg/kg, i.p.) was administered before behavioral testing. (**B**) Representative histological verification of viral expression in CA3. hM4Di-mCherry expression was predominantly localized to the deep CA3 pyramidal cell layer. (**C-E**) Open-field assessment of locomotor and anxiety-related behavior following CNO administration. (**C**) Schematic diagram for open-field test. (**D**) Time spent in the center. (**E**) Total distance traveled (*n* = 9-13 mice for each group). Unpaired *t* tests. (F-G) Contextual fear-memory retrieval following chemogenetic suppression of CA3 St18 neurons. (**F**) Schematic diagram of experimental procedure of contextual fear conditioning. Memory retrieval was tested 24 h (recent memory) and 21 days (remote memory) after foot shocks, with CNO administered 60 min before retrieval. (G) Freezing behavior in mCherry and hM4Di mice at either 24 h or 21 days. Two-way ANOVA retrieval time (F(1,30) = 27.93, *p* < 0.0001), group (F(1,30) = 0.024, *p* = 0.88) or group × retrieval-time interaction (F(1,30) = 0.37, *p* = 0.55). Sidak-adjusted comparisons showed no difference between mCherry and hM4Di mice at either the 24-h retrieval test (*p* = 0.83) or the 21-day remote-memory test (*p* = 0.94). (**H-J**) Y-maze spontaneous alternation test. (H) Schematic of the Y-maze paradigm. (I) Total arm entries. (J) Chemogenetic suppression of CA3 St18 neurons significantly reduced spontaneous alternation. *n* = 12-18 mice for each group. (K-L) T-maze spontaneous choice test. (**K**) Schematic of the spontaneous T-maze paradigm. (L) Correct-choice rate. Unpaired *t* tests, *n* = 12-13 mice per group. Data are presented as mean ± SEM. \**p* < 0.05, \*\**p* < 0.01. NS, not significant.

We then tested whether St18 neurons contribute to spatial working memory. In the Y-maze spontaneous alternation task (Fig. 7H), the number of arm entries was comparable between groups (t = 1.57, *p* = 0.13; Fig. 7I), indicating no changes in exploratory activity. By contrast, chemogenetic suppression of St18 neurons significantly reduced spontaneous alternation (mCherry, 62.2 ± 1.79%; hM4Di, 53.8 ± 3.77%; t = 2.21, *p* = 0.035; *n* = 12-18 mice per group; Fig. 7J). We independently assessed spatial working memory using a T-maze spontaneous choice task. Consistent with the Y-maze results, hM4Di mice showed a significantly lower correct choice rate than controls following CNO administration (mCherry, 79.5 ± 2.78%; hM4Di, 61.1 ± 5.60%; t = 3.00, *p* = 0.006; *n* = 12-13 mice per group; Fig. 7K-L). Together, these results support that St18 neuron activity contribute to performance in spontaneous spatial working memory tasks.

## DISCUSSION

We identify St18 neurons as a genetically accessible CA3 neuronal population with properties resembling previously described deep-layer, burst-firing athorny cells^2,26,42,43,46,47^, consistent with a recent study^50^. We further reveal that St18 neurons possess distinct features that distinguish them from canonical CA3 pyramidal neurons. St18 neuron outputs contrast sharply with canonical CA3 pyramidal neurons, with their axonal projections largely confined within CA3 recurrent network and only sparsely innervating CA1. St18 neurons strongly excite CA3 local interneurons, providing a potential mechanism for regulating recurrent excitation. St18 neurons are highly active during spontaneous CA3 SWR events, and their activation is sufficient to evoke ripple-like activity within CA3 but not CA1, whereas suppression reduces spontaneous CA3 ripple occurrence without affecting CA1. These region-specific effects are consistent with their preferred anatomical connections within CA3 and indicate that St18 neurons preferentially regulate local CA3 network dynamics. Finally, suppression of St18 neurons selectively impaired spontaneous spatial working memory without affecting contextual fear memory and other behavioral functions. Together, these findings identify St18 neurons as a molecularly defined CA3 circuit that has distinct intrinsic and connectivity features and contributes to both recurrent network dynamics and spatial working memory.

Although athorny cells have previously been described as a distinct CA3 subtype with unique morphophysiological properties^2,26,43^, their molecular identity has only recently been established. Consistent with a recent study^50^, our work demonstrates that St18 is a molecular marker of athorny cells. St18 neurons possess the cellular features defining athorny cells, including deep-layer position, burst-firing, and paucity of thorny excrescences. Thus, our St18-Cre line provides a genetic strategy to investigate this population beyond the morphological and physiological approaches used previously^2,26,42–44,47^. Interestingly, our transcriptomic analysis indicates that St18 neurons can be further resolved into multiple subclusters, highlighting the potential for functional diversity within St18 neurons.

Athorny neurons have previously been implicated in promoting synchronized firing in CA3 during SWR events, based on correlational analysis and computational modeling^2,26^. Yet, whether they are causally involved in these network dynamics is unknown. Our combined results using optotagged in vivo recordings and activity manipulations provide evidence that St18 neuron activity contributes to CA3 ripple dynamics. By contrast, we did not detect a significant change in CA1 ripple occurrence following St18 manipulation, consistent with their sparse direct projections to CA1. These findings support a preferential contribution of a molecularly defined CA3 population to local ripple dynamics, but do not exclude additional effects on downstream CA1 activity.

Our behavioral results suggest that St18 neurons contribute to rapid spatial updating. St18 neuron suppression impaired spatial working memory performance in two spatial working memory assays, whereas recognition memory and contextual fear memory were not affected. Together with the reduction in CA3 ripple occurrence, these findings support the view that St18-dependent CA3 network dynamics contribute to rapid updating of spatial information. Future work combining cell-type-specific manipulation with measurements of spatial representations and ripple-associated neuronal activity may clarify how St18-dependent network dynamics contribute to spatial working memory.

Overall, our findings suggest that the recurrent CA3 network contains molecularly defined neuronal populations with distinct circuit architecture and differential contributions to information flow through the trisynaptic circuit. St18 neurons are preferentially integrated into the recurrent CA3 network and selectively regulate CA3 ripple dynamics, with sparse direct output to CA1. The circuit architecture defined by St18 neurons provides a framework for understanding how CA3 cellular heterogeneity can shape hippocampal population dynamics and behavior.

## Acknowledgments

This work was supported by R01MH129294 and R01MH130367 from NIH in the United States (Q.S.) and institutional support from Case Western Reserve University (Q.S.). We thank the members in the Sun lab for helpful discussion.

## Author contributions

Conceptualization: WC, QS

Methodology: WC, HL, WG, DYW, EJB, YQJ, QS

Investigation: WC, HL, WG, DYW, EJB, YQJ, QS

Visualization: WC, HL, DYW, EJB, YQJ, QS

Funding acquisition: QS

Supervision: QS

Writing – original draft: WC

Writing – review & editing: QS

## Competing interests

Authors declare that they have no competing interests.

## Materials and Methods

### Mice

Experiments involving mice were approved by the Institutional Animal Care and Use Committee of Case Western Reserve University. Mice were housed on a 12-h light/dark cycle with free access to food and water. Wild-type (WT, stock No. 000664) and Grik4-Cre (stock No. 006474) mice were originally purchased from Jackson Laboratory. Mice used in all experiments were on a C57BL/6 background. St18-P2A-iCre (St18-Cre) knock-in mice were generated by Biocytogen (Waltham, MA) using the CRISPR-Cas9 knock-in strategy. To enable Cre recombinase expression under the control of the endogenous St18 regulatory elements while minimizing disruption of St18 expression, a P2A-iCre cassette was inserted immediately downstream of the coding sequence of exon 24 and upstream of the 3’ untranslated region (3’UTR) of the *St18* gene. The targeting construct contained approximately 1.5-kb 5’ and 3’ homology arms flanking the insertion site to facilitate homology-directed integration. St18-Cre knock-in mice exhibited no overt developmental abnormalities, gross alterations in brain morphology, or noticeable changes in body weight. Heterozygous St18-P2A-iCre mice of both sexes were used in the experiments. Genotyping of St18-P2A-iCre mice was performed using the following primers: WT-forward, 5′-GTT GGC AAC ATT TGT CAC TAT CTC C-3′; WT-reverse, 5′-TCC AGA AAG ATA TGT GAT TTG AGA ACA GC-3′; and mutant-reverse, 5′-CAG AGG GGC ATC CAC AGG GAG-3′. The WT allele yielded a 653-bp PCR product, whereas the mutant allele yielded a 370-bp product. All mouse lines were maintained on a C57BL/6 background.

### Brain slice electrophysiology

Mice were anesthetized with isoflurane and decapitated in accordance with institutional regulations. Brains were rapidly removed. Hippocampi were dissected in ice-cold dissection solution containing (in mM): 10 NaCl, 195 sucrose, 2.5 KCl, 10 glucose, 25 NaHCO₃, 1.25 NaH₂PO₄, 2 Na-Pyruvate, 0.5 CaCl₂, and 7 MgCl₂. The hippocampi were embedded in a block of pre-prepared 3–4% agarose and sectioned using a vibratome (Leica VT1000S or VT1200S) in ice-cold dissection solution. Transverse hippocampal slices (400 μm thick) were cut along the dorsal–ventral axis and transferred to artificial cerebrospinal fluid (ACSF) maintained at 32°C and containing (in mM): 125 NaCl, 2.5 KCl, 20 glucose, 25 NaHCO₃, 1.25 NaH₂PO₄, 2 Na-Pyruvate, 2 CaCl₂, and 1 MgCl₂. Slices were incubated in this solution for 20–30 min and then transferred to room temperature for at least 1.5 h before recording. Both the dissection solution and ACSF were continuously bubbled with 95% O₂ and 5% CO₂ to maintain a pH of 7.4. Electrophysiological recordings were performed at 31–32°C.

Whole-cell recordings were obtained from CA2 and CA3 pyramidal neurons, using both “visualized” and “blind” patch-clamp techniques under 40x or 4x objectives, as described previously^23,24,42,73^. Patch pipettes were pulled from borosilicate glass capillaries using a P-1000 micropipette puller (Sutter Instruments) and had resistances of approximately 4-6 MΩ. Pipettes were filled with an intracellular solution containing (in mM): 135 K-gluconate, 5 KCl, 0.1 Na-EGTA, 10 HEPES, 2 NaCl, 5 ATP, 0.4 GTP, and 10 phosphocreatine (pH 7.3-7.4; 280-290 mOsm). Biocytin (0.2-0.5%) was included in the intracellular solution for post hoc histological identification. Resting membrane potential was measured immediately after establishing whole-cell access. Neurons with resting membrane potentials more depolarized than −55 mV at break-in or series resistance greater than 30 MΩ were excluded from analysis. Liquid junction potentials were not corrected.

For voltage-clamp recordings, CA1 and CA3 neurons were held at −70 mV. Blue light pulses (470 nm, 2 ms) were delivered at maximum intensity using an LED light source (Thorlabs, LED4D294) controlled by pClamp 11 software and coupled to a 40× water-immersion objective positioned above the slice, as described previously^24,42,74^. Stimulation consisted of either a single light pulse or a train of pulses delivered at 20 Hz. To measure I/E ratio, cesium based intracellular solutions were used. The following drugs were bath-applied in ACSF as required: DNQX (20 μM) and DL-APV (50 μM).

For current-clamp recordings, postsynaptic potentials and action potentials were measured at the resting membrane potential. Frequency–current (F–I) curves were generated by injecting a series of current steps in 100-pA increments. Input resistance was calculated as the ratio of the steady-state change in membrane potential to the current amplitude during a 1-s, −100-pA current injection.

### Biocytin staining

Following whole-cell recordings, slices were fixed in 4% paraformaldehyde (PFA) for 24-48 h at 4°C. Fixed slices were incubated in blocking solution (PBS containing 5% normal goat serum and 0.5% Triton X-100) for 2 h at room temperature, followed by incubation with Streptavidin Alexa Fluor 594 (1:500; Invitrogen, S11227) or Streptavidin Alexa Fluor 647 (1:500; Invitrogen, S32357) in incubation solution (PBS containing 0.1% Triton X-100) for 2 days at 4°C. Slices were then washed three times in PBS for 10 min per wash and incubated with DAPI (0.5 μg/ml; Invitrogen, D1306) for 20–30 min at room temperature. After three additional PBS washes, slices were mounted on slides using VECTASHIELD Antifade Mounting Medium (Vector Laboratories, H-1900). Images were acquired using a confocal microscope (Zeiss LSM 800).

### Immunofluorescence

Mice were deeply anesthetized, perfused with phosphate buffered saline (PBS), followed by 4% paraformaldehyde (PFA). For coronal sections, whole brains were removed and post-fixed with 4% PFA overnight at 4°C. 50-60-µm sections were prepared on a vibratome (Leica VT1000S). Free-floating whole brain sections were processed for immunofluorescence. After three 10 min washes in PBS, sections were incubated with blocking solution (PBS with 5% normal goat serum and 0.3% triton X-100) for 2 hours. Then sections were incubated in primary antibodies in incubating solution (PBS with 3% normal goat serum and 0.3% triton X-100) for 1-2 days at 4℃. The following primary antibodies were used: chicken polyclonal anti-GFP (1:500, aves Labs, GFP-1020), and rat monoclonal anti-mCherry (1:500, Invitrogen, M11217). After being rinsed three times, each time for 10 min in PBS, sections were moved to secondary antibodies in incubating solution for 2 hours at room temperature. The following secondary antibodies were used: goat anti-chicken Alexa Fluor 488 (1:1000, Invitrogen, A32931TR) and goat anti-rat Alexa Fluor 594 (1:1000, Jackson ImmunoResearch, 112-585-167). After three 10 min washes in PBS, sections were incubated with DAPI (0.5 µg/ml, Invitrogen, D1306) for 20-30 min at room temperature. Then sections were rinsed in PBS for 3 × 10 min, before they were mounted with VECTASHIELD antifade mounting medium (Vector Laboratories, H-1000-10) on slides. The fluorescence images were obtained on a confocal microscope (Zeiss, LSM 800).

### Quantification of thorny excrescences

Coronal sections (50 µm) containing CA3 neurons labeled by AAV-DIO-EYFP injections in Grik4-Cre and St18-Cre mice were immunostained with an anti-GFP antibody to enhance the visualization of neuronal morphology. High-resolution confocal z-stack images were acquired using a Zeiss LSM microscope equipped with Airyscan and imported into Imaris 9.0 (Bitplane, US) for 3D reconstruction. Dendritic shafts and associated protrusions, including conventional spines and simple and complex thorny excrescences, were reconstructed using the Filament Tracer module. Reconstructions were initiated at the proximal portion of the dendrites and extended along the dendritic branches toward stratum lucidum and stratum radiatum. Dendritic shafts and their associated spines and thorny excrescences were traced separately. The volumes of individual dendritic segments and their associated protrusions were obtained from the reconstructed structures. For each neuron, total dendritic shaft volume was calculated by summing the volumes of all reconstructed dendritic segments, explicitly excluding the volumes of spines and thorny excrescences. Total protrusion volume was calculated as the sum of the volumes of all conventional spines and simple and complex thorny excrescences associated with the reconstructed dendrites. To account for differences in dendritic size and branching complexity, protrusion volume was normalized to the corresponding dendritic shaft volume for each neuron. The resulting dimensionless ratio, calculated as total protrusion volume divided by total dendritic shaft volume, was used to compare dendritic protrusion morphology between Grik4-Cre- and St18-Cre-labeled CA3 pyramidal neurons.

### Neuronal reconstruction and analysis

Imaris software (Oxford Instruments) was used to analyze dendritic length (apical and basal), soma volume, and Sholl intersections. Z-stack images of biocytin-filled neurons were captured using a Zeiss LSM 800 confocal microscope with a 20x objective. The images were converted into Imaris files and rendered in 3D. Dendrites were traced using the *Filament* module with the semi-automatic *AutoPath* algorithm (minimum diameter set to 0.60 µm), with all paths originating from the soma center. Apical and basal dendrites were traced as separate filament objects. The soma was reconstructed using the *Surface* module by defining a region of interest around the cell body and applying the automatic creation wizard. Relevant morphological parameters were then extracted from the Imaris statistics tab.

### Single-cell RNA-seq analysis

#### Data source and dataset selection

Single-cell RNA-seq data were obtained from the Allen Institute cortex and hippocampal formation atlas (GEO:GSE185862)^69^. Data from adult mouse isocortex and hippocampal formation were profiled using both 10x Genomics Chromium v2 and SMART-seq v4 platforms. Brain-region boundaries were defined according to the Allen Mouse Brain Common Coordinate Framework version 3 (CCFv3), and hippocampal formation dissections included the tissue containing CA3. Publicly available human single-nucleus RNA-seq data were obtained from the Allen Brain Cell Atlas (WHB-10Xv3 dataset)^70^. Nuclei from the hippocampal CA3 region were selected based on anatomical annotations and analyzed to characterize transcriptional heterogeneity and gene expression patterns across CA3 neuronal populations.

#### Extraction of CA3 neuronal transcriptomes

Metadata and expression matrices for both 10x and SMART-seq datasets were processed in R. Cells were retained CA3-related anatomical annotations. Neuronal cells of CA3 were further restricted to annotated glutamatergic neurons. Cell identifiers were harmonized between metadata and expression matrices by testing barcode-format transformations and selecting the mapping with maximal overlap.

#### Integration and downstream analysis

CA3 neurons from the 10x and SMART-seq datasets were integrated using Seurat^75^. Genes shared across platforms were retained, mitochondrial transcript percentage was calculated, and each dataset was independently normalized using SCTransform. Integration features were selected, SCT integration was performed using reciprocal PCA anchors, and the integrated object was subjected to PCA, UMAP visualization, nearest-neighbor graph construction, and graph-based clustering.

#### Differential gene expression and identification of domain-enriched markers

Differential gene expressions between CA3 domains were evaluated using Seurat’s FindMarkers function with a Wilcoxon rank-sum test. The right domain (clusters 1, 2, 3, 5, 6, and 7) was designated as ident.1, and the left domain (clusters 0 and 4) as ident.2; therefore, positive log2 fold-change values indicated higher expression in the right domain. Candidate domain-enriched markers were identified using predefined criteria: Bonferroni-adjusted P < 0.05, expression fold change > 3, detection in > 30% of cells in the target domain, and detection in < 15% of cells in every cluster belonging to the opposite domain. In addition, each cluster within the target domain was required to exhibit a detection rate > 30%. Identical criteria were applied to identify genes enriched in either the left or right domain. Genes that failed to meet any of these criteria were not classified as domain-enriched markers. The number and percentage of cells in each category were quantified separately for each cluster and for the pooled left and right domains.

### RNAscope in situ hybridization and Quantification

#### RNAscope

RNAscope in situ hybridization was performed using the RNAscope Multiplex Fluorescent Reagent Kit v2 (Cat. No. 323100; ACD, USA) according to the manufacturer’s instructions. Briefly, mouse brains were fixed in 4% paraformaldehyde and cryoprotected in a graded sucrose series (20-30%). Coronal sections (15 µm) containing the hippocampal CA3 region were prepared using a cryostat. Sections were subsequently subjected to pretreatment, target retrieval, and permeabilization using Manual PretreatPro reagents (ACD). RNAscope probes targeting mouse *St18* (Mm-St18-C3; Cat. No. 443271-C3) and *P2rx7* (Mm-P2rx7-C2; Cat. No. 316311-C2) were obtained from ACD. Following probe hybridization and signal amplification, *St18* and *P2rx7* transcripts were visualized using Opal 520 and Opal 690 fluorophores and followed by immunofluorescence staining.

#### Image processing and nuclear segmentation

Regions of interest (ROIs) encompassing hippocampal CA3 were manually delineated, and the deep and superficial laminar boundaries were traced based on DAPI staining. Nuclei were segmented within the CA3 ROIs using a threshold-based watershed approach. Small foreground objects were removed, and internal holes were filled. Nuclear labels were subsequently expanded by 5-10 pixels to generate cell-associated measurement regions for RNA signal quantification.

#### RNA signal quantification

Fluorescence signals corresponding to *P2rx7* and *St18* transcripts were background subtracted. Pixels exceeding the intensity threshold were identified as candidate RNA puncta and assigned to the nearest expanded nuclear measurement region. *St18* RNA signals formed dense, spatially overlapping clusters that prevented reliable identification of individual puncta, the primary quantitative measure was the area occupied by positive fluorescence.

#### Laminar localization and depth-dependent expression analysis

To characterize the laminar distribution of *P2rx7* and *St18* expression within hippocampal CA3, the shortest distances from each nuclear centroid to the manually delineated deep and superficial boundaries were calculated. The normalized depth axis was divided into 30 equally spaced bins. Mean RNA signal was calculated for each image and subsequently averaged within each field group to generate depth-dependent expression profiles. For the normalized overlay visualization, each field-group profile was standardized by z-score normalization. The final depth profiles are presented as the mean ± standard error of the mean. Negative z-scores indicate RNA signal levels below the mean of the corresponding field-group profile.

### Surgery and viral injections

For in vivo experiments, AAVs expressing GFP, mCherry, or ChR2 were diluted two-fold (1:2, ∼1.0-2.0 ×10^12^ GC/ml), whereas AAV-hM4Di was administered without dilution. AAV injection surgeries were performed following protocols from previous studies^76^. Briefly, mice were anesthetized with 2.5% isoflurane in a 30% O₂: 70% N₂ gas mixture and placed on a heating pad to maintain body temperature at 37°C. The animals were then secured in a stereotactic apparatus for stereotaxic surgery. Holes targeting the CA3 (AP: −2.0 mm, ML: 2.90 mm, DV: 2.4 mm) were drilled, and AAVs were delivered into the target brain region via a fine glass electrode connected to an infusion pump, with an infusion rate of 0.05 µl/min (KDS101, KD Scientific). Afterward, the skin was sutured, and the animals were allowed to recover for two weeks. Subsequently, the mice were subjected to behavioral and electrophysiological experiments.

### In vivo electrophysiology

#### Surgery and electrophysiological recordings

In vivo electrophysiological recordings were performed in St18-Cre mice (3-5-month-old) expressing mCherry, hM4Di, or ChR2 in hippocampal CA3 St18 neurons. Mice were initially anesthetized with isoflurane (2.5% in a 30%:70% O2:N2 gas mixture), and anesthesia was maintained at lighter level (0.7-0.8%) throughout the recording session. Animals were secured in a stereotaxic apparatus, and lidocaine was administered subcutaneously at the incision site for local anesthesia. Following a scalp incision, small craniotomies were made over hippocampal CA1b (AP: −2.1 mm; ML: 1.9 mm; DV: 1.2-1.4 mm) and CA3 (AP: −2.1 mm; ML: 2.9 mm; DV: 1.6-2.3 mm). Body temperature was maintained at ∼37°C throughout the electrophysiological recording session. All stereotaxic coordinates were referenced to bregma, with dorsoventral coordinates measured from. Carbon fiber microelectrodes (7 µm in diameter) were fabricated as previously described^76,77^ and trimmed to expose approximately 50 µm of the fiber. The electrodes had an impedance of approximately 5 MΩ. Carbon fiber electrodes were positioned in CA1b or CA3 for extracellular electrophysiological recordings. For optogenetic experiments, an optical fiber (200 µm diameter) was positioned above CA3 (AP: −2.1 mm; ML: 3.3 mm; DV: 1.6 mm) using a micromanipulator tilted at 20° relative to the vertical axis. A reference electrode was positioned over the cerebellum and secured using an anchoring screw. Extracellular signals were initially amplified using a 10× preamplifier, followed by additional filtering and amplification (gain: 200) using a differential amplifier (EX1; Dagan Corporation, USA). Signals were digitized using a National Instruments data acquisition card (NI PCIe-6351; National Instruments, USA) and acquired using WaveSurfer software. Following electrophysiological recordings, mice were euthanized, and brain tissue was collected for histological verification of electrode placement and viral expression.

#### Data analysis and spike sorting

Extracellular recordings from hippocampal CA3 were acquired from multiple mice at a sampling rate of 20 kHz. Spike sorting was performed using MountainSort5 through the SpikeInterface framework^78,79^. Sorting results were exported to Phy2 for manual inspection and curation^80^. Clusters were visually inspected and classified with manual merging or splitting. Spike trains from labeled clusters were retained for subsequent analyses, except for units that were manually excluded at the analysis stage according to predefined exclusion criteria.

#### Spike-sorting quality control

Spike-sorting quality was assessed retrospectively using archived Phy2 outputs. Quality metrics included unit presence ratio, refractory-period violations (interspike intervals <1 and < 2 ms), spike-amplitude variability, amplitude-time correlation, and autocorrelograms.

#### Optogenetic identification of putative St18 neurons

Putative St18 neurons were identified using an optogenetic tagging approach adapted from previously established methods with slight modification^81^. Extracellular single-unit activity was recorded from hippocampal CA3 using carbon fiber microelectrodes while ChR2-expressing St18 neurons were stimulated with blue light (473 nm). Optical stimulation consisted of 100-ms light pulses (∼2.4 mW measured at the fiber tip), delivered over 60-120 trials. Stimulation was controlled using WaveSurfer software through the analog output of a National Instruments data acquisition device connected to a BNC-2110 (NI) terminal block. Putative St18 neurons were identified based on three criteria: (1) a median first-spike latency of less than 7 ms following light onset; (2) a first-spike response reliability of at least 60% across stimulation trials^82^; and (3) an increase in firing rate during the light pulse, including a sustained response during the late stimulation phase (50-100 ms after light onset). Only units meeting all three criteria were included in subsequent analyses of putative St18 neurons. Units that failed to meet these criteria were excluded from the optogenetically identified St18 population.

#### Complex-spike detection and characterization

Complex-spike bursts were detected from spike trains recorded during spontaneous activity epochs. Burst detection was based on interspike interval (ISI) criteria, consistent with established approaches for characterizing hippocampal pyramidal-cell bursting^83^. A primary burst event was defined as a maximal, nonoverlapping sequence of at least three consecutive spikes, with all ISIs ≤ 20 ms. Each event was time-stamped at the onset of its first spike. Burst frequency was calculated as the number of detected burst onsets divided by the total duration of eligible recording epochs. Burst size was calculated as the mean number of spikes per event, and intraburst timing was characterized by the median ISI within each burst. Bursts exhibiting a progressive decrease in positive spike amplitude across successive spikes were identified as amplitude-decrementing events.

#### Classification of theta-dominant and non-theta-dominant activity

Theta and delta power were calculated over 5-12 Hz and 1-4 Hz, respectively. To account for unequal bandwidths, classification was based on the ratio of mean PSD: 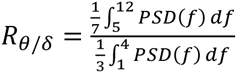 Each spectral estimate was assigned to a nonoverlapping 1 s interval centered on the analysis window and was considered scoreable only when the entire contributing 2 s window contained valid data. In the primary analysis, ratios ≥ 1.5 were classified as theta-dominant, ratios ≤ 0.8 as non-theta-dominant, and intermediate values as uncertain. Same-state periods shorter than 3 s were reassigned as uncertain, followed by exclusion of 1 s margins around state transitions. States were not extended across sweep boundaries, artifacts, or acquisition gaps.

#### Ripple detection, induction and ripple-associated firing

Spontaneous ripples were detected from the continuous recording using criteria and MATLAB code from Buzsaki lab with slight modification^55,58,84^. Briefly, wideband signals (0.1-10 kHz) were low-pass filtered and downsampled to 1,250 Hz to obtain LFPs, while signals high-pass filtered above 0.5 kHz were used for spike detection. Ripple detection parameters were adjusted to retain events lasting 20-120 ms. Recording sites in CA1b and CA3 were verified post hoc by histological examination. Animals initially received an intraperitoneal injection of saline to establish baseline activity. Then, CNO (2 mg/kg, i.p.) was administered, and spontaneous ripples were recorded and quantified beginning 45 mins after injection. We further analyzed the recording sessions by separating them into theta and non-theta epochs (see above). Only non-theta epochs were included in the subsequent analysis of spontaneous ripple activity.

#### Optogenetics

In the optogenetic induction of ripple-like event, a ramp stimulation waveform using WaveSurfur and analog output devices (BNC-2110, NI).^55^ The optogenetic stimulation, lasting 100 ms were applied to the CA3 with the laser power tunned from 0.7 to 3 mW (MBL-F-473-100 mW, PSU-H-LED, Cnlaser, China) to minimize the stimulation artifacts.

#### Association of spontaneous ripple-peak

Spikes and complex-spike event onsets were aligned to ripple peaks from −200 to +200 ms in 10-ms bins. For each neuron, firing rates were z-scored using the combined flanking baseline (−200 to −50 ms and +50 to +200 ms). Peak activity was defined as the mean z-score within ± 25 ms of the ripple peak. Units with insufficient data or zero baseline variance were excluded. Population activity is presented as mean ± SEM across neurons. Complex-spike event rates were z-scored against the combined flanking baseline (−200 to −50 ms and +50 to +200 ms). Peak complex-spike activity was defined as the mean z-score within ± 25 ms of the ripple peak.

#### Sharp-wave detection and association

Sharp waves (SWs) were detected independently of ripple oscillations using a modified approach based on previous study with slight modification^26^. Raw electrophysiological signals were low-pass filtered at 100 Hz, downsampled to 1 kHz, and subsequently band-pass filtered at 2-50 Hz using a zero-phase filter. The filtered signal was normalized by subtracting the median and dividing by 1.48 times the median absolute deviation (MAD). Candidate SW events were detected when the absolute normalized signal reached or exceeded a threshold of 3, with event boundaries defined by an absolute z-score of 1.5. Events separated by ≤ 20 ms were merged, and only events lasting 25-200 ms were retained. Each event was aligned to the time of its largest absolute LFP deflection, and events were pooled for subsequent analyses regardless of polarity.

### Behavioral Tests

Behavioral experiments were performed using 3-5-month-old mice. Mice were handled and habituated for at least 3 consecutive days (10 min/mice/day) before behavioral testing. Health and condition of the mice were examined routinely. Experimenters were blind to the manipulations as well as genotypes. On each testing day, mice were allowed to acclimate to the behavioral testing room or apparatus before the start of the experiment. Mice expressing hM4Di and control mice expressing mCherry received CNO (2.0 mg/kg, i.p.) Both hM4Di and control mice received CNO injections 1 h before behavioral testing.

#### T-maze spontaneous alternation

Spontaneous alternation was assessed using a T-maze as previously described with slight modification^85,86^. Each mouse underwent seven consecutive trials, consisting of an initial sample trial followed by six test trials. Both goal arms remained accessible throughout testing. Following each choice, mice were confined to the selected arm for 30 s before being returned to the start arm. An alternation was defined as entry into the arm opposite to that selected in the preceding trial. Spontaneous alternation was expressed as the percentage of correct alternations across the six test trials.

#### Y-maze spontaneous alternation

Spatial working memory was assessed using a Y-maze spontaneous alternation task. Mice were placed individually in the maze and allowed to freely explore all three arms for 10 min. Arm-entry sequences and total entries were recorded, with an entry defined as all four paws entering an arm. An alternation was defined as consecutive entries into three different arms. Spontaneous alternation was calculated as the number of actual alternations divided by the total possible alternations (total arm entries - 2), expressed as a percentage.

#### Open-field test

Locomotor activity and exploratory behaviors were assessed in an open-field arena. Mice were placed individually in the arena and allowed to explore freely for 30 min. The center zone was defined as the central 25% of the total arena. Total distance traveled, and time spent in the center was quantified.

#### Contextual fear conditioning

Contextual fear memory was assessed using a standard fear conditioning paradigm. Following an initial habituation period, mice received four foot-shocks (1 s each, 0.5 mA), separated by intertrial intervals. Contextual memory retrieval was assessed 24 h after conditioning by re-exposing mice to the original conditioning chamber without shock delivery. Remote memory retrieval was performed 20 days after conditioning using the same procedure. Freezing behavior, defined as the absence of movement except that required for respiration, was quantified as the percentage of time spent freezing during each retrieval session.

#### Novel object recognition

Recognition memory was assessed using the novel object recognition task. Mice were first habituated to an empty arena for 10 min. During familiarization, two identical objects were placed in the arena, and mice were allowed to explore for 10 min. After a 24-h retention interval, one familiar object was replaced with a novel object, and exploratory behavior was recorded for 10 min. Object exploration was defined as active investigation directed toward an object, excluding climbing or incidental contact. Recognition performance was quantified using the discrimination index: (novel object exploration time − familiar object exploration time)/(total object exploration time).

### Statistical analysis

Statistical analyses were performed using GraphPad Prism 10.0 (GraphPad Software, USA) or OriginPro 2022 (OriginLab, Northampton, MA, USA). Data are presented as mean ± SEM unless otherwise indicated. Normality was assessed using the Shapiro-Wilk test. Comparisons between two independent groups were performed using unpaired *t* tests. Experiments involving two independent factors were analyzed using Two-way ANOVA, or repeated-measures two-way ANOVA when measurements were obtained from the same animals, followed by Sidak’s and Tukey’s multiple-comparisons. Differential gene-expression analyses were performed using the Wilcoxon rank-sum test with correction for multiple testing. All statistical tests were two-sided, and statistical significance was defined as *p* < 0.05.

## Notes

### Competing Interest Statement

The authors have declared no competing interest.

## References

1. Kong, E., Zabeh, E., Liao, Z., Mihaila, T.S., Wilson, C., Santhirasegaran, C., Peterka, D.S., Geiller, T., and Losonczy, A. (2026). Recurrent connectivity shapes spatial coding in hippocampal CA3 subregions. Neuron. 10.1016/j.neuron.2026.06.010.

2. Sammons, R.P., Masserini, S., Moreno Velasquez, L., Metodieva, V.D., Cano, G., Sannio, A., Orlando, M., Maier, N., Kempter, R., and Schmitz, D. (2025). Sub-type specific connectivity between CA3 pyramidal neurons may underlie their sequential activation during sharp waves. Elife 13. 10.7554/eLife.98653.

3. Layous, R., Tamas, B., Mike, A., Sipos, E., Arszovszki, A., Brunner, J., Szatai, A., Yaseen, F., Andrasi, T., and Szabadics, J. (2025). Optical Recordings of Unitary Synaptic Connections Reveal High and Random Local Connectivity between CA3 Pyramidal Cells. J Neurosci 45. 10.1523/JNEUROSCI.0102-25.2025.

4. Rebola, N., Carta, M., and Mulle, C. (2017). Operation and plasticity of hippocampal CA3 circuits: implications for memory encoding. Nat Rev Neurosci 18, 208–220. 10.1038/nrn.2017.10.

5. Guzman, S.J., Schlogl, A., Frotscher, M., and Jonas, P. (2016). Synaptic mechanisms of pattern completion in the hippocampal CA3 network. Science 353, 1117–1123. 10.1126/science.aaf1836.

6. Le Duigou, C., Simonnet, J., Telenczuk, M.T., Fricker, D., and Miles, R. (2014). Recurrent synapses and circuits in the CA3 region of the hippocampus: an associative network. Front Cell Neurosci 7, 262. 10.3389/fncel.2013.00262.

7. Nakazawa, K., Quirk, M.C., Chitwood, R.A., Watanabe, M., Yeckel, M.F., Sun, L.D., Kato, A., Carr, C.A., Johnston, D., Wilson, M.A., and Tonegawa, S. (2002). Requirement for hippocampal CA3 NMDA receptors in associative memory recall. Science 297, 211–218. 10.1126/science.1071795.

8. Debanne, D., Gahwiler, B.H., and Thompson, S.M. (1998). Long-term synaptic plasticity between pairs of individual CA3 pyramidal cells in rat hippocampal slice cultures. J Physiol 507 (Pt 1), 237–247. 10.1111/j.1469-7793.1998.237bu.x.

9. Miles, R., and Wong, R.K. (1986). Excitatory synaptic interactions between CA3 neurones in the guinea-pig hippocampus. J Physiol 373, 397–418. 10.1113/jphysiol.1986.sp016055.

10. Witter, M.P. (2007). Intrinsic and extrinsic wiring of CA3: indications for connectional heterogeneity. Learn Mem 14, 705–713. 10.1101/lm.725207.

11. Watson, J.F., Vargas-Barroso, V., Morse-Mora, R.J., Navas-Olive, A., Tavakoli, M.R., Danzl, J.G., Tomschik, M., Rossler, K., and Jonas, P. (2024). Human hippocampal CA3 uses specific functional connectivity rules for efficient associative memory. Cell. 10.1016/j.cell.2024.11.022.

12. Rolls, E.T. (2013). A quantitative theory of the functions of the hippocampal CA3 network in memory. Front Cell Neurosci 7, 98. 10.3389/fncel.2013.00098.

13. McHugh, T.J., and Tonegawa, S. (2009). CA3 NMDA receptors are required for the rapid formation of a salient contextual representation. Hippocampus 19, 1153–1158. 10.1002/hipo.20684.

14. Nakashiba, T., Buhl, D.L., McHugh, T.J., and Tonegawa, S. (2009). Hippocampal CA3 output is crucial for ripple-associated reactivation and consolidation of memory. Neuron 62, 781–787. 10.1016/j.neuron.2009.05.013.

15. Nakashiba, T., Young, J.Z., McHugh, T.J., Buhl, D.L., and Tonegawa, S. (2008). Transgenic inhibition of synaptic transmission reveals role of CA3 output in hippocampal learning. Science 319, 1260–1264. 10.1126/science.1151120.

16. Nakazawa, K., Sun, L.D., Quirk, M.C., Rondi-Reig, L., Wilson, M.A., and Tonegawa, S. (2003). Hippocampal CA3 NMDA receptors are crucial for memory acquisition of one-time experience. Neuron 38, 305–315.

17. Robert, V., O’Neil, K., Moore, J.J., Rashid, S.K., Johnson, C.D., De La Torre, R.G., Zemelman, B.V., Clopath, C., and Basu, J. (2025). Cortical glutamatergic and GABAergic inputs support learning-driven hippocampal stability. Science 390, eadn0623. 10.1126/science.adn0623.

18. Li, Y., Briguglio, J.J., Romani, S., and Magee, J.C. (2024). Mechanisms of memory-supporting neuronal dynamics in hippocampal area CA3. Cell 187, 6804–6819 e6821. 10.1016/j.cell.2024.09.041.

19. Pelkey, K.A., Vargish, G.A., Pellegrini, L.V., Calvigioni, D., Chapeton, J., Yuan, X., Hunt, S., Cummins, A.C., Eldridge, M.A.G., Pickel, J., et al. (2023). Evolutionary conservation of hippocampal mossy fiber synapse properties. Neuron 111, 3802–3818 e3805. 10.1016/j.neuron.2023.09.005.

20. Middleton, S.J., and McHugh, T.J. (2016). Silencing CA3 disrupts temporal coding in the CA1 ensemble. Nat Neurosci 19, 945–951. 10.1038/nn.4311.

21. Vyleta, N.P., Borges-Merjane, C., and Jonas, P. (2016). Plasticity-dependent, full detonation at hippocampal mossy fiber-CA3 pyramidal neuron synapses. Elife 5. 10.7554/eLife.17977.

22. Li, M., Jiang, Y.Q., Lee, D.K., Wang, H., Lu, M.C., and Sun, Q. (2024). Dorsoventral Heterogeneity of Synaptic Connectivity in Hippocampal CA3 Pyramidal Neurons. J Neurosci 44. 10.1523/JNEUROSCI.0370-24.2024.

23. Sun, Q., Jiang, Y.Q., and Lu, M.C. (2020). Topographic heterogeneity of intrinsic excitability in mouse hippocampal CA3 pyramidal neurons. J Neurophysiol 124, 1270–1284. 10.1152/jn.00147.2020.

24. Sun, Q., Sotayo, A., Cazzulino, A.S., Snyder, A.M., Denny, C.A., and Siegelbaum, S.A. (2017). Proximodistal Heterogeneity of Hippocampal CA3 Pyramidal Neuron Intrinsic Properties, Connectivity, and Reactivation during Memory Recall. Neuron 95, 656–672 e653. 10.1016/j.neuron.2017.07.012.

25. Thompson, C.L., Pathak, S.D., Jeromin, A., Ng, L.L., MacPherson, C.R., Mortrud, M.T., Cusick, A., Riley, Z.L., Sunkin, S.M., Bernard, A., et al. (2008). Genomic anatomy of the hippocampus. Neuron 60, 1010–1021. 10.1016/j.neuron.2008.12.008.

26. Hunt, D.L., Linaro, D., Si, B., Romani, S., and Spruston, N. (2018). A novel pyramidal cell type promotes sharp-wave synchronization in the hippocampus. Nat Neurosci 21, 985–995. 10.1038/s41593-018-0172-7.

27. Cembrowski, M.S., and Spruston, N. (2019). Heterogeneity within classical cell types is the rule: lessons from hippocampal pyramidal neurons. Nat Rev Neurosci 20, 193–204. 10.1038/s41583-019-0125-5.

28. Cembrowski, M.S., Wang, L., Sugino, K., Shields, B.C., and Spruston, N. (2016). Hipposeq: a comprehensive RNA-seq database of gene expression in hippocampal principal neurons. Elife 5, e14997. 10.7554/eLife.14997.

29. Ishizuka, N., Cowan, W.M., and Amaral, D.G. (1995). A quantitative analysis of the dendritic organization of pyramidal cells in the rat hippocampus. J Comp Neurol 362, 17–45. 10.1002/cne.903620103.

30. Ishizuka, N., Weber, J., and Amaral, D.G. (1990). Organization of intrahippocampal projections originating from CA3 pyramidal cells in the rat. J Comp Neurol 295, 580–623. 10.1002/cne.902950407.

31. Kjelstrup, K.B., Solstad, T., Brun, V.H., Hafting, T., Leutgeb, S., Witter, M.P., Moser, E.I., and Moser, M.B. (2008). Finite scale of spatial representation in the hippocampus. Science 321, 140–143. 10.1126/science.1157086.

32. Lu, L., Igarashi, K.M., Witter, M.P., Moser, E.I., and Moser, M.B. (2015). Topography of Place Maps along the CA3-to-CA2 Axis of the Hippocampus. Neuron 87, 1078–1092. 10.1016/j.neuron.2015.07.007.

33. Lee, H., Wang, C., Deshmukh, S.S., and Knierim, J.J. (2015). Neural Population Evidence of Functional Heterogeneity along the CA3 Transverse Axis: Pattern Completion versus Pattern Separation. Neuron 87, 1093–1105. 10.1016/j.neuron.2015.07.012.

34. Li, X.G., Somogyi, P., Ylinen, A., and Buzsaki, G. (1994). The hippocampal CA3 network: an in vivo intracellular labeling study. J Comp Neurol 339, 181–208. 10.1002/cne.903390204.

35. Kowalski, J., Gan, J., Jonas, P., and Pernia-Andrade, A.J. (2016). Intrinsic membrane properties determine hippocampal differential firing pattern in vivo in anesthetized rats. Hippocampus 26, 668–682. 10.1002/hipo.22550.

36. Oliva, A., Fernandez-Ruiz, A., Buzsaki, G., and Berenyi, A. (2016). Spatial coding and physiological properties of hippocampal neurons in the Cornu Ammonis subregions. Hippocampus 26, 1593–1607. 10.1002/hipo.22659.

37. Hunsaker, M.R., Rosenberg, J.S., and Kesner, R.P. (2008). The role of the dentate gyrus, CA3a,b, and CA3c for detecting spatial and environmental novelty. Hippocampus 18, 1064-1073. 10.1002/hipo.20464.

38. Kesner, R.P. (2007). Behavioral functions of the CA3 subregion of the hippocampus. Learn Mem 14, 771–781. 10.1101/lm.688207.

39. Lee, H., GoodSmith, D., and Knierim, J.J. (2020). Parallel processing streams in the hippocampus. Curr Opin Neurobiol 64, 127–134. 10.1016/j.conb.2020.03.004.

40. Turner, D.A., Li, X.G., Pyapali, G.K., Ylinen, A., and Buzsaki, G. (1995). Morphometric and electrical properties of reconstructed hippocampal CA3 neurons recorded in vivo. J Comp Neurol 356, 580–594. 10.1002/cne.903560408.

41. Bilkey, D.K., and Schwartzkroin, P.A. (1990). Variation in electrophysiology and morphology of hippocampal CA3 pyramidal cells. Brain Res 514, 77–83. 10.1016/0006-8993(90)90437-g.

42. Li, M., Jiang, Y.Q., and Sun, Q. (2025). A circuit from the basolateral amygdala to hippocampal CA3 regulates social behavior. Curr Biol 35, 4349–4364 e4345. 10.1016/j.cub.2025.07.059.

43. Watson, J.F., Vargas-Barroso, V., and Jonas, P. (2025). Cell-specific wiring routes information flow through hippocampal CA3. Cell Rep 44, 116080. 10.1016/j.celrep.2025.116080.

44. Raus Balind, S., Mago, A., Ahmadi, M., Kis, N., Varga-Nemeth, Z., Lorincz, A., and Makara, J.K. (2019). Diverse synaptic and dendritic mechanisms of complex spike burst generation in hippocampal CA3 pyramidal cells. Nat Commun 10, 1859. 10.1038/s41467-019-09767-w.

45. Marissal, T., Bonifazi, P., Picardo, M.A., Nardou, R., Petit, L.F., Baude, A., Fishell, G.J., Ben-Ari, Y., and Cossart, R. (2012). Pioneer glutamatergic cells develop into a morpho-functionally distinct population in the juvenile CA3 hippocampus. Nat Commun 3, 1316. 10.1038/ncomms2318.

46. Linaro, D., Levy, M.J., and Hunt, D.L. (2022). Cell type-specific mechanisms of information transfer in data-driven biophysical models of hippocampal CA3 principal neurons. PLoS Comput Biol 18, e1010071. 10.1371/journal.pcbi.1010071.

47. Kis, N., Luko, B., Heredi, J., Mago, A., Erlinghagen, B., Ahmadi, M., Raus Balind, S., Iras, M., Ujfalussy, B.B., and Makara, J.K. (2024). Cholinergic regulation of dendritic Ca(2+) spikes controls firing mode of hippocampal CA3 pyramidal neurons. Proc Natl Acad Sci U S A 121, e2321501121. 10.1073/pnas.2321501121.

48. Zheng, Z., Park, C., Hammerschmith, E.W., Lu, R., Yu, S.C., Sorek, M., Silverman, B., Jordan, C.S., Sterling, A.R., Silversmith, W.M., et al. (2026). Hippocampal CA3 connectomics reveals a gradient of mossy fiber inputs and selective feedforward inhibition onto pyramidal cells. Nat Neurosci 29, 2311–2323. 10.1038/s41593-026-02388-9.

49. Balleza-Tapia, H., Arroyo-Garcia, L.E., Isla, A.G., Loera-Valencia, R., and Fisahn, A. (2022). Functionally-distinct pyramidal cell subpopulations during gamma oscillations in mouse hippocampal area CA3. Prog Neurobiol 210, 102213. 10.1016/j.pneurobio.2021.102213.

50. Morse-Mora, R.J., Ben-Simon, Y., Arcot Jayaram, S., Rössler, K., Jonas, P., and Watson, J.F. (2026). Hippocampal CA3 forms a two-layer network of molecularly distinct cell types in mice and humans. bioRxiv.

51. Buzsaki, G. (2015). Hippocampal sharp wave-ripple: A cognitive biomarker for episodic memory and planning. Hippocampus 25, 1073–1188. 10.1002/hipo.22488.

52. Colgin, L.L. (2016). Rhythms of the hippocampal network. Nat Rev Neurosci 17, 239–249. 10.1038/nrn.2016.21.

53. Roumis, D.K., and Frank, L.M. (2015). Hippocampal sharp-wave ripples in waking and sleeping states. Curr Opin Neurobiol 35, 6–12. 10.1016/j.conb.2015.05.001.

54. Buzsaki, G. (1986). Hippocampal sharp waves: their origin and significance. Brain Res 398, 242–252. 10.1016/0006-8993(86)91483-6.

55. Fernandez-Ruiz, A., Oliva, A., Fermino de Oliveira, E., Rocha-Almeida, F., Tingley, D., and Buzsaki, G. (2019). Long-duration hippocampal sharp wave ripples improve memory. Science 364, 1082–1086. 10.1126/science.aax0758.

56. Csicsvari, J., Hirase, H., Mamiya, A., and Buzsaki, G. (2000). Ensemble patterns of hippocampal CA3-CA1 neurons during sharp wave-associated population events. Neuron 28, 585–594. 10.1016/s0896-6273(00)00135-5.

57. Jadhav, S.P., Kemere, C., German, P.W., and Frank, L.M. (2012). Awake hippocampal sharp-wave ripples support spatial memory. Science 336, 1454–1458. 10.1126/science.1217230.

58. Oliva, A., Fernandez-Ruiz, A., Buzsaki, G., and Berenyi, A. (2016). Role of Hippocampal CA2 Region in Triggering Sharp-Wave Ripples. Neuron 91, 1342–1355. 10.1016/j.neuron.2016.08.008.

59. Girardeau, G., Benchenane, K., Wiener, S.I., Buzsaki, G., and Zugaro, M.B. (2009). Selective suppression of hippocampal ripples impairs spatial memory. Nat Neurosci 12, 1222–1223. 10.1038/nn.2384.

60. Singer, A.C., Carr, M.F., Karlsson, M.P., and Frank, L.M. (2013). Hippocampal SWR activity predicts correct decisions during the initial learning of an alternation task. Neuron 77, 1163–1173. 10.1016/j.neuron.2013.01.027.

61. Papale, A.E., Zielinski, M.C., Frank, L.M., Jadhav, S.P., and Redish, A.D. (2016). Interplay between Hippocampal Sharp-Wave-Ripple Events and Vicarious Trial and Error Behaviors in Decision Making. Neuron 92, 975–982. 10.1016/j.neuron.2016.10.028.

62. Joo, H.R., and Frank, L.M. (2018). The hippocampal sharp wave-ripple in memory retrieval for immediate use and consolidation. Nat Rev Neurosci 19, 744–757. 10.1038/s41583-018-0077-1.

63. Foster, D.J., and Wilson, M.A. (2006). Reverse replay of behavioural sequences in hippocampal place cells during the awake state. Nature 440, 680–683. 10.1038/nature04587.

64. Csicsvari, J., and Dupret, D. (2014). Sharp wave/ripple network oscillations and learning-associated hippocampal maps. Philos Trans R Soc Lond B Biol Sci 369, 20120528. 10.1098/rstb.2012.0528.

65. Hwaun, E., and Colgin, L.L. (2019). CA3 place cells that represent a novel waking experience are preferentially reactivated during sharp wave-ripples in subsequent sleep. Hippocampus. 10.1002/hipo.23090.

66. Bazelot, M., Telenczuk, M.T., and Miles, R. (2016). Single CA3 pyramidal cells trigger sharp waves in vitro by exciting interneurones. J Physiol 594, 2565–2577. 10.1113/JP271644.

67. Schlingloff, D., Kali, S., Freund, T.F., Hajos, N., and Gulyas, A.I. (2014). Mechanisms of sharp wave initiation and ripple generation. J Neurosci 34, 11385–11398. 10.1523/JNEUROSCI.0867-14.2014.

68. Hajos, N., Karlocai, M.R., Nemeth, B., Ulbert, I., Monyer, H., Szabo, G., Erdelyi, F., Freund, T.F., and Gulyas, A.I. (2013). Input-output features of anatomically identified CA3 neurons during hippocampal sharp wave/ripple oscillation in vitro. J Neurosci 33, 11677–11691. 10.1523/JNEUROSCI.5729-12.2013.

69. Yao, Z., van Velthoven, C.T.J., Nguyen, T.N., Goldy, J., Sedeno-Cortes, A.E., Baftizadeh, F., Bertagnolli, D., Casper, T., Chiang, M., Crichton, K., et al. (2021). A taxonomy of transcriptomic cell types across the isocortex and hippocampal formation. Cell 184, 3222–3241 e3226. 10.1016/j.cell.2021.04.021.

70. Siletti, K., Hodge, R., Mossi Albiach, A., Lee, K.W., Ding, S.L., Hu, L., Lonnerberg, P., Bakken, T., Casper, T., Clark, M., et al. (2023). Transcriptomic diversity of cell types across the adult human brain. Science 382, eadd7046. 10.1126/science.add7046.

71. Stark, E., Roux, L., Eichler, R., Senzai, Y., Royer, S., and Buzsaki, G. (2014). Pyramidal cell-interneuron interactions underlie hippocampal ripple oscillations. Neuron 83, 467–480. 10.1016/j.neuron.2014.06.023.

72. Oliva, A., Fernandez-Ruiz, A., Leroy, F., and Siegelbaum, S.A. (2020). Hippocampal CA2 sharp-wave ripples reactivate and promote social memory. Nature 587, 264–269. 10.1038/s41586-020-2758-y.

73. Sun, Q., Srinivas, K.V., Sotayo, A., and Siegelbaum, S.A. (2014). Dendritic Na(+) spikes enable cortical input to drive action potential output from hippocampal CA2 pyramidal neurons. Elife 3. 10.7554/eLife.04551.

74. Kinney, J.L., Robinson, H.L., Wang, H., Jiang, Y.Q., Zhou, M., Lee, D.K., Li, M., Oliva, A., and Sun, Q. (2025). Top-down regulation of subcortical regions by hippocampal long-range inhibition. Curr Biol 35, 5853–5867 e5854. 10.1016/j.cub.2025.10.062.

75. Hao, Y., Stuart, T., Kowalski, M.H., Choudhary, S., Hoffman, P., Hartman, A., Srivastava, A., Molla, G., Madad, S., Fernandez-Granda, C., and Satija, R. (2024). Dictionary learning for integrative, multimodal and scalable single-cell analysis. Nat Biotechnol 42, 293–304. 10.1038/s41587-023-01767-y.

76. Cui, W., Gao, N., Dong, Z., Shen, C., Zhang, H., Luo, B., Chen, P., Comoletti, D., Jing, H., Wang, H., et al. (2021). In trans neuregulin3-Caspr3 interaction controls DA axonal bassoon cluster development. Curr Biol 31, 3330–3342 e3337. 10.1016/j.cub.2021.05.045.

77. Fortin, S.M., Cone, J.J., Ng-Evans, S., McCutcheon, J.E., and Roitman, M.F. (2015). Sampling phasic dopamine signaling with fast-scan cyclic voltammetry in awake, behaving rats. Curr Protoc Neurosci 70, 7 25 21–27 25 20. 10.1002/0471142301.ns0725s70.

78. Buccino, A.P., Hurwitz, C.L., Garcia, S., Magland, J., Siegle, J.H., Hurwitz, R., and Hennig, M.H. (2020). SpikeInterface, a unified framework for spike sorting. Elife 9. 10.7554/eLife.61834.

79. Chung, J.E., Magland, J.F., Barnett, A.H., Tolosa, V.M., Tooker, A.C., Lee, K.Y., Shah, K.G., Felix, S.H., Frank, L.M., and Greengard, L.F. (2017). A Fully Automated Approach to Spike Sorting. Neuron 95, 1381–1394 e1386. 10.1016/j.neuron.2017.08.030.

80. Rossant, C., Kadir, S.N., Goodman, D.F.M., Schulman, J., Hunter, M.L.D., Saleem, A.B., Grosmark, A., Belluscio, M., Denfield, G.H., Ecker, A.S., et al. (2016). Spike sorting for large, dense electrode arrays. Nat Neurosci 19, 634–641. 10.1038/nn.4268.

81. Lima, S.Q., Hromadka, T., Znamenskiy, P., and Zador, A.M. (2009). PINP: a new method of tagging neuronal populations for identification during in vivo electrophysiological recording. PLoS One 4, e6099. 10.1371/journal.pone.0006099.

82. Lakunina, A.A., Socha, K.Z., Ladd, A.E., Bowen, A.J., Chen, S., Colonell, J., Doshi, A., Karsh, B., Krumin, M., Kulik, P., et al. (2026). Neuropixels Opto: combining high-resolution electrophysiology and optogenetics. Nat Methods 23, 1207–1216. 10.1038/s41592-026-03076-z.

83. Harris, K.D., Hirase, H., Leinekugel, X., Henze, D.A., and Buzsáki, G. (2001). Temporal interaction between single spikes and complex spike bursts in hippocampal pyramidal cells. Neuron 32, 141–149. Doi 10.1016/S0896-6273(01)00447-0.

84. Liu, A.A., Henin, S., Abbaspoor, S., Bragin, A., Buffalo, E.A., Farrell, J.S., Foster, D.J., Frank, L.M., Gedankien, T., Gotman, J., et al. (2022). A consensus statement on detection of hippocampal sharp wave ripples and differentiation from other fast oscillations. Nat Commun 13, 6000. 10.1038/s41467-022-33536-x.

85. Deacon, R.M., and Rawlins, J.N. (2006). T-maze alternation in the rodent. Nat Protoc 1, 7–12. 10.1038/nprot.2006.2.

86. d’Isa, R., Comi, G., and Leocani, L. (2021). Apparatus design and behavioural testing protocol for the evaluation of spatial working memory in mice through the spontaneous alternation T-maze. Sci Rep-Uk 11. ARTN 21177 10.1038/s41598-021-00402-7.

