## Supplemental Figures for "A molecularly defined recurrent CA3 circuit regulates network dynamics and spatial working memory"

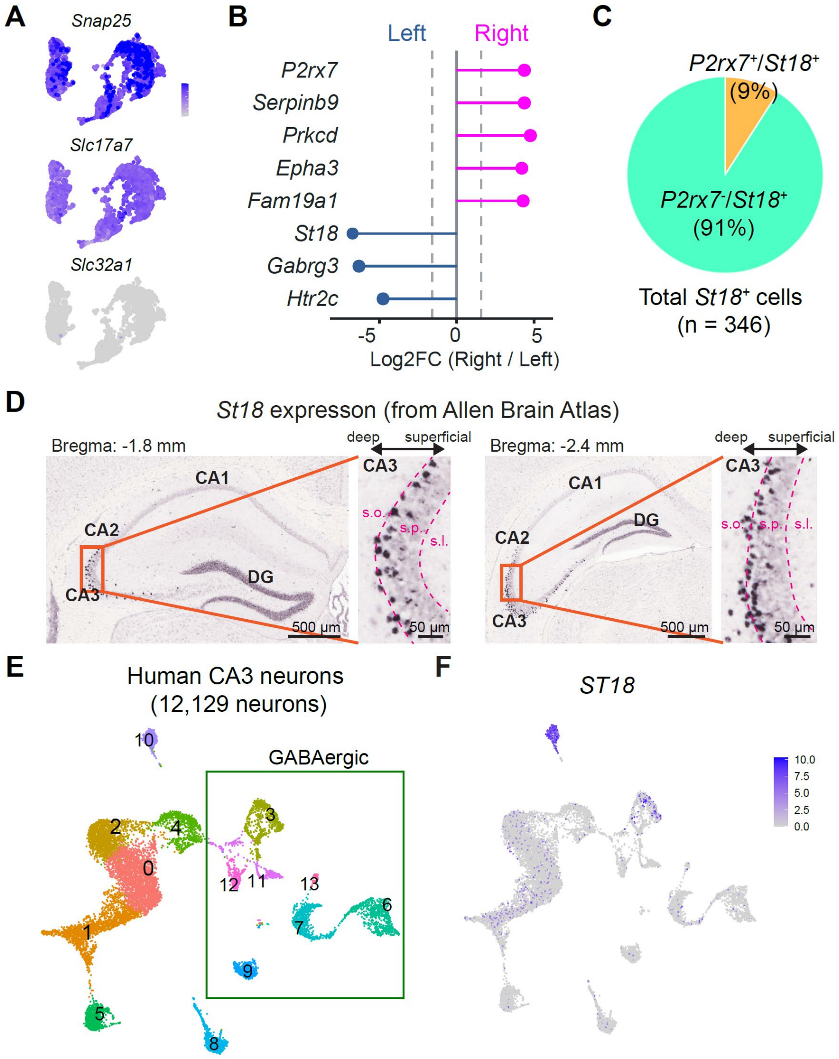


**Figure S1. Single-cell transcriptome analysis in mouse and human CA3.**
(A) UMAP feature plots showing expression of the pan-neuronal marker *Snap25*, glutamatergic marker *Slc17a7/Vglut1* and GABAergic neuron marker *Slc32a1* in CA3 neurons.
(B) Fold-enrichment analysis of candidate marker genes in the left and right CA3 transcriptional domains.
(C) Colocalization of *St18* and *P2rx7* expression in mouse CA3. *n* = 346 cells from 2 mice.
(D) Representative in situ hybridization images showing *St18* expression in dorsal and ventral CA3 obtained from the Allen Brain Atlas.
(E-F) Analysis of *ST18* expression in the human hippocampal CA3. (D) UMAP representation of human CA3 neuronal populations showing glutamatergic and GABAergic neuronal clusters. (E) UMAP feature plot showing *ST18*. Scale, [0 10].

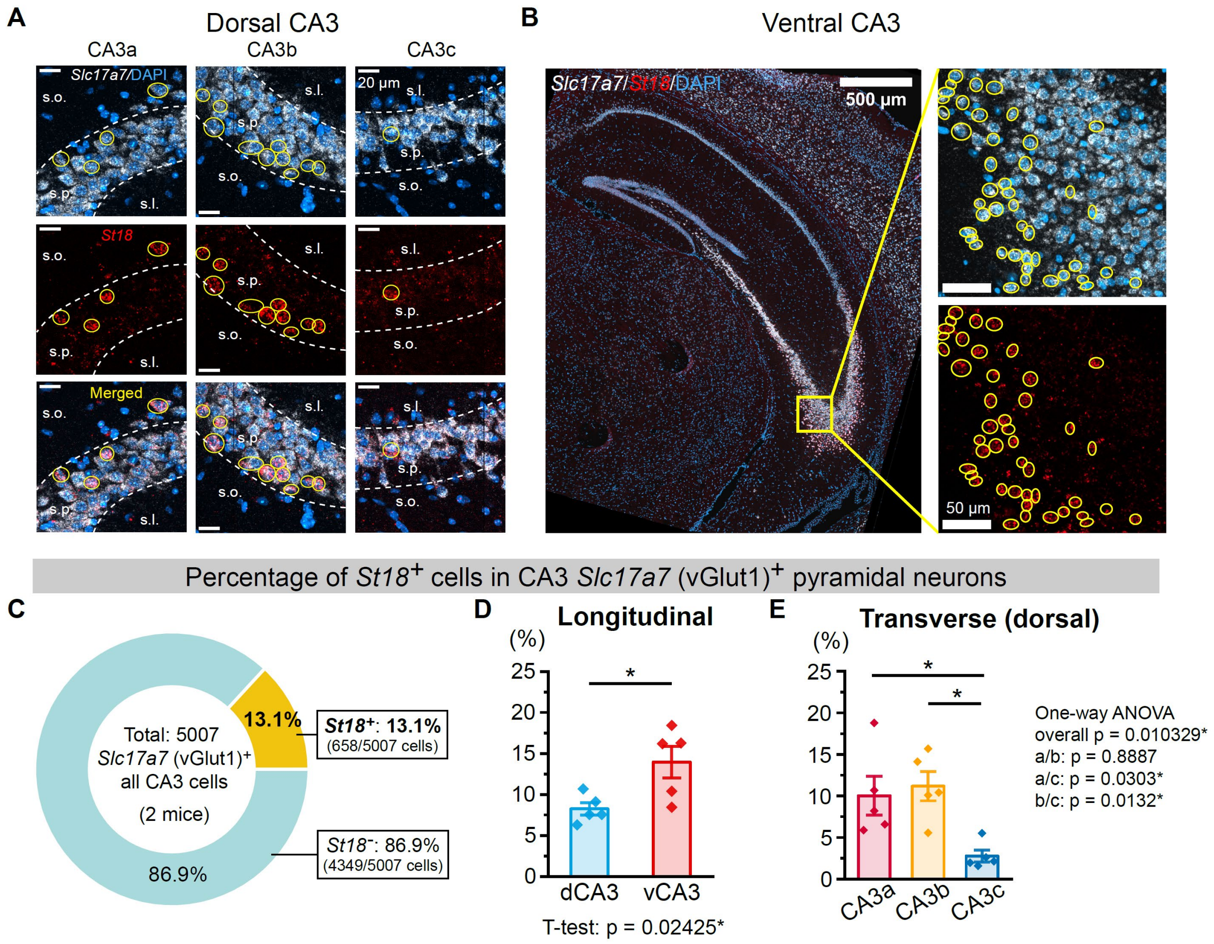


**Figure S2. Distribution of St18 neurons along the transverse and longitudinal axes.**

(A) Representative confocal images of RNA fluorescent in situ hybridization (RNAscope) for *St18* and *Slc17a7* (vGlut1) along the transverse axis of dorsal CA3. Open yellow circles indicate *St18*$^+$/*Slc17a7*$^+$ cells.

(B) Representative confocal images of RNA fluorescent in situ hybridization for *St18* and *Slc17a7* (vGlut1) in ventral CA3. Open yellow circles indicate *St18*$^+$/*Slc17a7*$^+$ cells.

(C) Quantification of the percentage of *Slc17a7*$^+$ cells expressing *St18* in CA3. $n$ = 5007 *Slc17a7*$^+$ cells from 2 mice.

(D) Quantification of the percentage of *Slc17a7*$^+$ cells expressing *St18* in dorsal CA3 (dCA3) and ventral CA3 (vCA3). $n$ = 5 hippocampi from 2 mice per group.

(E) Quantification of the percentage of *Slc17a7*$^+$ cells expressing *St18* along the transverse axis of dCA3. $n$ = 5 sections from 2 mice per group. Error bars represent SEM.

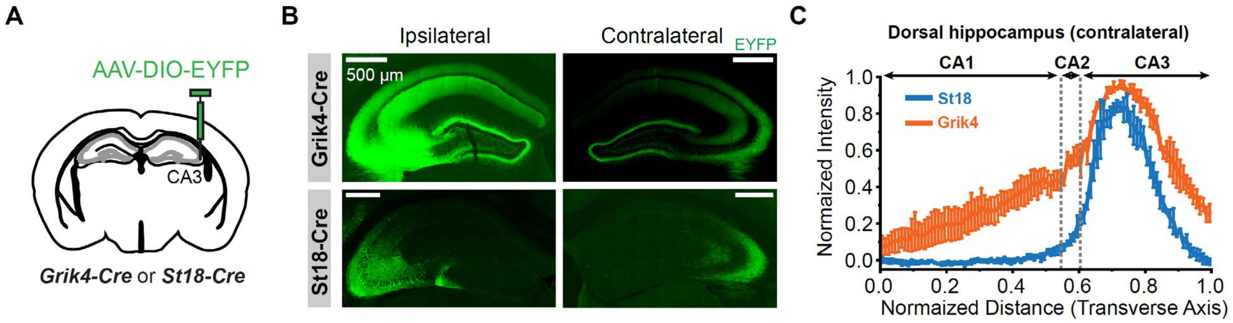


**Figure S3. Axonal projections of St18 neurons compared with the broader CA3 pyramidal neuron population.**
(A) AAV-DIO-EYFP was injected unilaterally into CA3 of Grik4-Cre or St18-Cre mice.
(B) Representative confocal images of coronal sections showing EYFP-labeled axonal projections in the ipsilateral and contralateral hippocampus from Grik4-Cre (top) and St18-Cre (bottom) mice.
(C) Quantification of normalized EYFP fluorescence intensity along the CA1-CA3 transverse axis in Grik4-Cre ($n$ = 5 sections/3 mice) and St18-Cre mice ($n$ = 6 sections/3 mice).

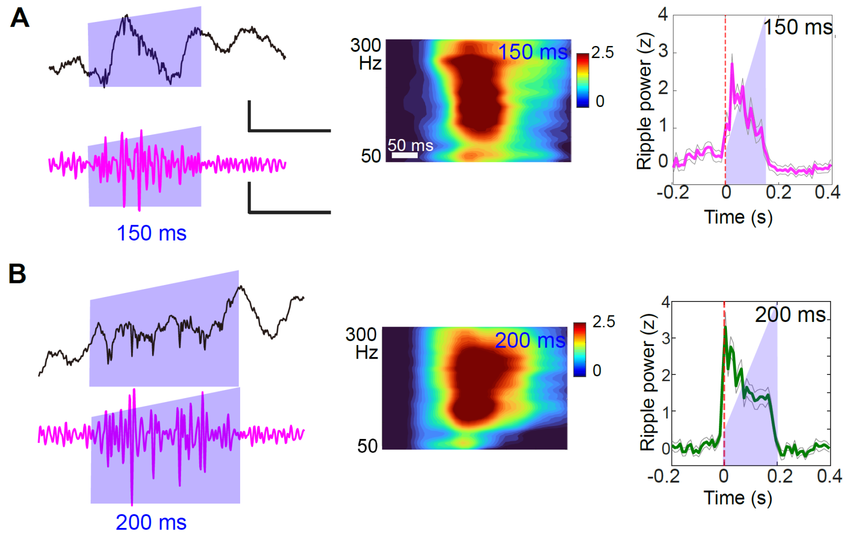


**Figure S4. Ripple-like high-frequency oscillations evoked by optogenetic activation of CA3 St18 neurons with different durations.**
(A-B) Representative CA3 LFP and ripple-band-filtered responses to prolonged optogenetic stimulation of St18 neurons. Stimulation durations of 150 ms (A) and 200 ms (B). Representative LFP, ripple band, time frequency decomposition and peri-stimulation ripple power were shown. Blue shading indicates the photo-stimulation period. Scale bars, 1 mV for LFP, 0.2 mV for ripple band, time 100 ms. Color scales, [0 2.5]. Time frequency decomposition, scale, 50 ms. *n* = 4 mice.

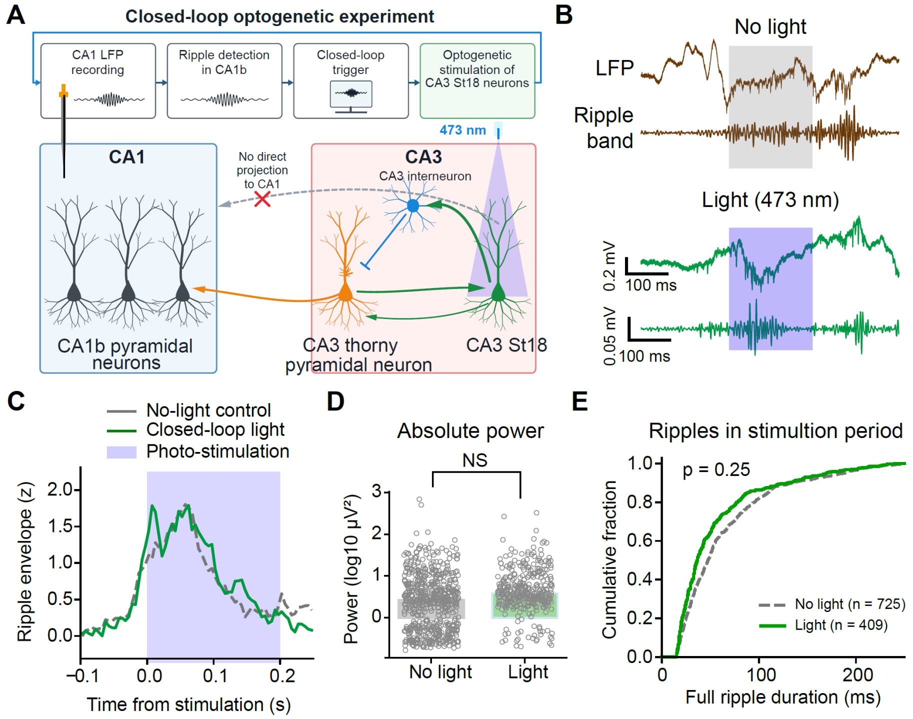


**Figure S5. Closed-loop St18 stimulation did not produce an increase in CA1 ripple power.**
(A) Schematic of CA1 ripple recording and closed-loop photo-stimulation.
(B) Representative traces of LFP and ripple band in no-light and light stimulation. Brown traces, no-light control, green, light stimulation.
(C) Mean z-scored ripple-band envelope aligned to photostimulation onset. Blue shading indicates the photostimulation window. Lines mean ripple power.
(D) Quantification of ripple power during the stimulation window under no-light control and photostimulation conditions. A total of 650 sham stimulation events and 360 photostimulation events were analyzed across 3 mice.
(E) Cumulative distribution of ripple durations during the stimulation period. *n* = 3 mice. Wilcoxon matched pairs signed rank test, W = 0, *p* = 0.25.

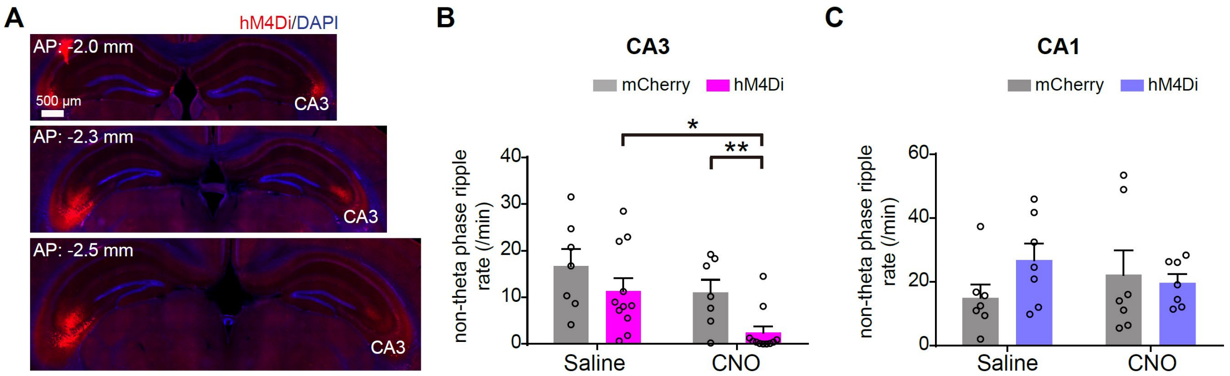


**Figure S6. Chemogenetic suppression of CA3 St18 neurons reduces CA3 ripple activity during non-theta periods.**
(A) Representative coronal brain sections illustrating bilateral hM4Di-mCherry expression in CA3 of St18-Cre mice following AAV injection, arranged along the anterior–posterior axis.
(B) Quantification of CA3 ripple occurrence rate during non-theta periods following saline or CNO administration in mCherry-expressing control mice and hM4Di-expressing mice. Two-way repeated-measures ANOVA revealed a significant main effect of treatment ($F(1,16) = 6.83$, $p = 0.018$). $n = 7$ mCherry control mice and 11 hM4Di mice.
(C) Quantification of CA1 ripple occurrence rate during non-theta periods following saline or CNO administration in mCherry-expressing control mice and hM4Di-expressing mice. CNO administration did not significantly affect CA1 ripple occurrence rate (main effect of treatment: $F(1,12) = 0.0002$, $p = 0.99$; saline versus CNO: mCherry, $p = 0.56$; hM4Di, $p = 0.57$). $n = 7$ mice per group.
Data are presented as mean ± SEM. Two-way repeated-measures ANOVA followed by Sidak's multiple-comparisons test. *$p < 0.05$; **$p < 0.01$.

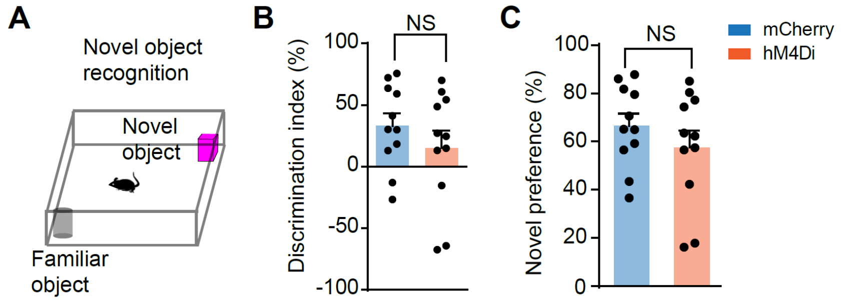


**Figure S7. Chemogenetic suppression of CA3 St18 neurons does not impair novel object recognition.**
(A-C) Novel object recognition test following chemogenetic suppression of CA3 St18 neurons. (A) Schematic of the novel object recognition paradigm. (B) Discrimination index. t = 1.0, *p* = 0.32. (C) Novel-object preference did not differ between mCherry and hM4Di mice following CNO administration (t = 1.00, *p* = 0.32). Unpaired *t* tests, *n* = 11 mice for each group.
